# Flexible Methods for Species Distribution Modeling with Small Samples

**DOI:** 10.1101/2025.04.03.647025

**Authors:** Brian S. Maitner, Robert L. Richards, Ben Carlson, John M. Drake, Cory Merow

## Abstract

Species distribution models (SDMs) are used for understanding where species live or could potentially live and are a key resource for ecological research and conservation decision-making. However, current SDM methods often perform poorly for rare or inadequately sampled species, which includes most species on earth as well as most of those of the greatest conservation concern.
Here, we evaluate the performance of three recently developed modeling approaches specifically designed for data-deficient situations: 1) plug-and-play modeling, 2) density-ratio modeling, and 3) environmental-range modeling. We compare the performance of these methods with Maxent, a widely used method. We compare model performance across sample sizes as well as comparisons limited to only data-poor species. We also ask to what extent model cross-validation performance on training data was correlated with model performance on independent, presence-absence data.
We show that, across all species, one or more of the plug-and-play, density-ratio, or environmental-range algorithms outperformed Maxent in 72% of cases, with three of the algorithms having AUC distributions not significantly different from Maxent’s. For data-poor species (those with 20 or fewer occurrences), 24 of the algorithms considered had AUC distributions that were not significantly different from Maxent. However, despite these comparable AUC scores, we found that the algorithm outputs (when thresholded to predict presence vs absence) spanned a wide gradient of sensitivity vs. specificity. Specificity and prediction accuracy assessed on training data were strongly correlated with specificity and prediction accuracy assessed on independent presence-absence data, however AUC and sensitivity had weak correlations. We found that only for 16% of species was the model that performed best on the training data the best performing model when evaluated on independent, presence-absence data. Finally, we show how ensembles of models that span the sensitivity-specificity gradient can represent model disagreement in poorly sampled species and improve model predictions.
This work supports plug-and-play, density-ratio, and environmental-range modeling as useful alternatives to Maxent, particularly for data-deficient species. While our work suggests that identifying the best model for a given species is challenging, we argue that incorporating the predictions of multiple models provides a useful way forward.

**Data/Code for peer review:** Anonymized data and code underlying this work are publicly available at: https://anonymous.4open.science/r/Flexible_Methods_for_Small_Sample_Size_Distribution_Modeling

## Introduction

Delineating locations where species occur or could potentially occur, *e.g.*, due to introductions or migration following climate change, is critically important for both advancing our understanding of global biodiversity and ensuring it is adequately protected. Species distribution models (SDMs) provide reproducible methods to identify the potential spatial distributions of species as a function of their environments (Peterson et al., 2011; S. J. Phillips et al., 2006). SDMs are widely used (Araújo et al., 2019) to understand where species are located (*e.g.*, B. S. Maitner et al., 2017), where they could be located after introduction or migration (*e.g.* Hannah et al., 2020), and to understand their ecology and evolution (*e.g.*, Neves et al., 2021). SDMs can be built using different types of data, including presence-only data (Busby, 1991; Drake, 2014; Drake & Richards, 2018), presence-background data (S. J. Phillips et al., 2006; Renner et al., 2015; Warton & Shepherd, 2010), presence-absence data (Thuiller et al., 2016), abundance data (Royle, 2004), or repeated measures of any of these data types (Royle & Dorazio, 2008). In this study we focus on the first two, as absence and abundance data are not available for most species.

With increasing recognition of the value of SDMs there has been an explosion of methodology for building them (reviewed in: Elith et al., 2006; Guillera-Arroita et al., 2015; Merow et al., 2014; Norberg et al., 2019; Valavi et al., 2022). Despite these advances, the ability to accurately model the distributions of species with small sample sizes (*e.g.*, rare, poorly sampled, or small-ranged species) remains a persistent problem (Breiner et al., 2018; Hernandez et al., 2006; Jeliazkov et al., 2022; Zhang et al., 2020). Across the globe, most species are rare (Enquist et al., 2019), and global sampling biases (Meyer et al., 2015) mean that only a few occurrence records may be available for sometimes wide-spread species (Hughes et al., 2021; B. Maitner et al., 2023; B. S. Maitner et al., 2017). For example, of an estimated 360,000 to 400,000 known plant species, around 293,000 (73 - 81 percent) have publicly-available, spatially-referenced occurrence data that pass data cleaning standards in the BIEN database (B. S. Maitner et al., 2017). Of these species with data, only approximately 112,000 (38 percent) have more than 10 unique records suitable for use with traditional range modeling methods. Thus there is a critical need for methods to estimate species distributions from only a handful of occurrence records--if that task can be done at all.

Small sample sizes are a ubiquitous problem for SDMs. While many algorithms have been shown to be accurate with sufficient data, few have been shown to consistently provide useful results with small sample sizes (Breiner et al., 2018; Wisz et al., 2008). While many algorithms do not converge with small sample sizes, some simpler algorithms that will converge, including BioClim (Busby, 1991), have been shown to perform poorly with small sample sizes (Hernandez et al., 2006). Other algorithms, including Maxent, have been shown to perform well with small sample sizes in special cases (Galante et al., 2018; Hernandez et al., 2006), but this is not a general result and may depend on having an unbiased sample (Jeliazkov et al., 2022). Unfortunately, sampling biases cannot be precisely assessed for poorly-sampled species represented only by a few presence records, and any assessment must rely on assumptions that are difficult to validate and may only provide imperfect corrections (Boyd et al., 2023, 2024). Hence to address bias researchers are forced to make assumptions to correct for it, and report uncertainty or ensemble results over different assumptions.

### Models for data-poor species

We focus on three modeling approaches that can be applied to data-poor species: 1) plug-and-play modeling, 2) density-ratio modeling, and 3) environmental-range modeling. The first of these approaches, plug-and-play modeling (Figure 1), integrates core ecological concepts, and allows for a flexible approach to distribution modeling (Drake & Richards, 2018). In the plug-and-play framework, the joint distribution of all environmental covariates of interest is defined as the environmental distribution, f(z), and the set of environmental covariates at which a species occurs is defined as its environmental distribution, f_1_(z). We refer to f(z) as the background distribution and f_1_(z) as the presence distribution. Drake and Richards show how the relative occurrence rate (ROR), *i.e.*, the relative probability that the sample derived from each cell in the domain (termed “relative suitability” in the framework of Drake & Richards, 2018; “relative occurrence rate” in Fithian & Hastie, 2013; Merow et al., 2013), S_R_(z) of a species can be estimated as S_R_(z) = f_1_(z)/f(z) (Fig. 1). Thus, SDMs can be constructed by “plugging in” any algorithm that can estimate these distributions. Alternatively, the second set of algorithms that we consider directly estimate the ratio of these density functions (rather than estimating the presence and background distributions separately). Density-ratio algorithms include unconstrained least-squares importance fitting (uLSIF) and relative unconstrained least-squares importance fitting (ruLSIF) (Drake & Richards, 2018; Kanamori et al., 2009; Yamada et al., 2013). Maxent (S. J. Phillips et al., 2006) models also fall into the class of density ratio models, as they estimate S_R_(z), while imposing additional constraints on model fitting to penalize for model complexity (Drake & Richards, 2018). The third modeling approach, environmental-range modeling, omits the use of the background distribution and instead focuses on identifying the environmental limits of a species’ niche; examples include range-bagging (Drake, 2015) and (low-bias bootstrap-aggregating, one-class; Drake, 2014), abbreviated LOBAG-OC (Drake, 2014).

**Figure 1.**
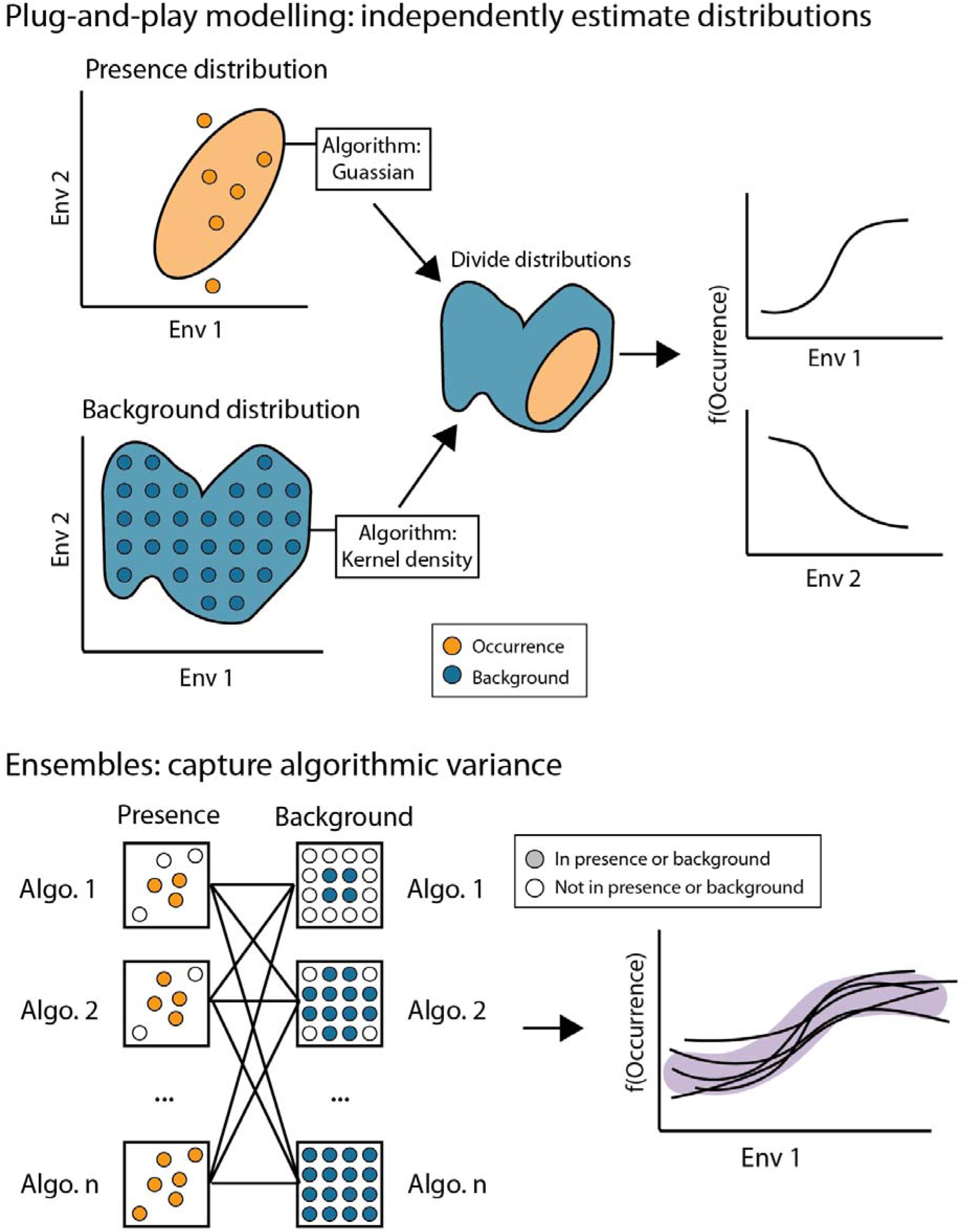
Using Plug-and-Play Modeling to capture uncertainty for small sample size species. The Plug-and-play framework estimates the relative occurrence rate, f(occurrence), as the ratio of a presence distribution and a background distribution (Upper panel). These distributions can be fit using different algorithms that reflect diverse assumptions about sampling decisions or use-cases. Alternatively, algorithms that can directly estimate the ratio of these two distributions can be used, reflecting their own assumptions. In most cases it will be unknown which assumptions are met. Creating an ensemble of multiple algorithms and combinations of algorithms representing a range of assumptions provides a way to capture this uncertainty (bottom panel), which is particularly important for small sample size species.

The flexibility of these three modeling approaches allows them to incorporate multiple algorithms, which we propose can be interpreted through their different assumptions about sampling bias and niche shape. These sampling assumptions can be conceptualized by thinking of algorithms along a sensitivity-specificity gradient (Figure 2). Sensitivity is the probability of correctly predicting presences, while specificity is the probability of correctly predicting absences. While the exact values of sensitivity and specificity will depend on the thresholds used to convert continuous model outputs to presences and absences, the relative positions along a sensitivity-specificity gradient will be consistent. On one end of the spectrum, Kernel Density Estimation (KDE) can achieve high specificity by providing a relatively tight fight to the observed data (depending on tuning parameters), and thus can reflect the assumption that the occurrence records accurately capture the environment a species occupies; KDE can also be tuned to generate high sensitivity, but less specific models by increasing the bandwidth. On the other end of the spectrum, Gaussian distributions and Maxent (optionally, depending on settings) achieve high sensitivity by providing a looser fit to the observed data (i.e., including presences that are further from known occurrences in environmental space), and are appropriate where we expect that there are likely to be suitable environments that are relatively distance from known occurrences in environmental space. Other algorithms may range between these extremes, providing different compromises between sensitivity and specificity. Importantly, there is no *a priori* reason why the same algorithm should be used for estimating *f_1_(**z**)* and *f(**z**)*, and we might expect better performance when we combine different algorithms. For example, where a species has few records we may want a relatively high sensitivity estimate of *f_1_(**z**)*, but we may have enough background records that we prefer a high specificity *f(**z**)*. These methods also differ in the assumptions they make about niche shape. For example, range-bagging focuses on estimating the boundaries of a distribution, and hence isn’t strongly affected by sampling bias within distributions. Alternatively, KDE distributions allow for discontinuous niches while gaussian distributions do not. Since these methods can capture a wide set of assumptions, we hypothesized that they may be particularly useful for very small sample sizes (*i.e.*, 1 - 20 records) where uncertainty is important and assumptions are difficult to validate. We also propose that ensembles of the algorithms presented here, that differ in their assumptions, can more precisely characterize ranges, along with their uncertainty, in spite of small sample sizes and in the presence of bias.

**Figure 2.**
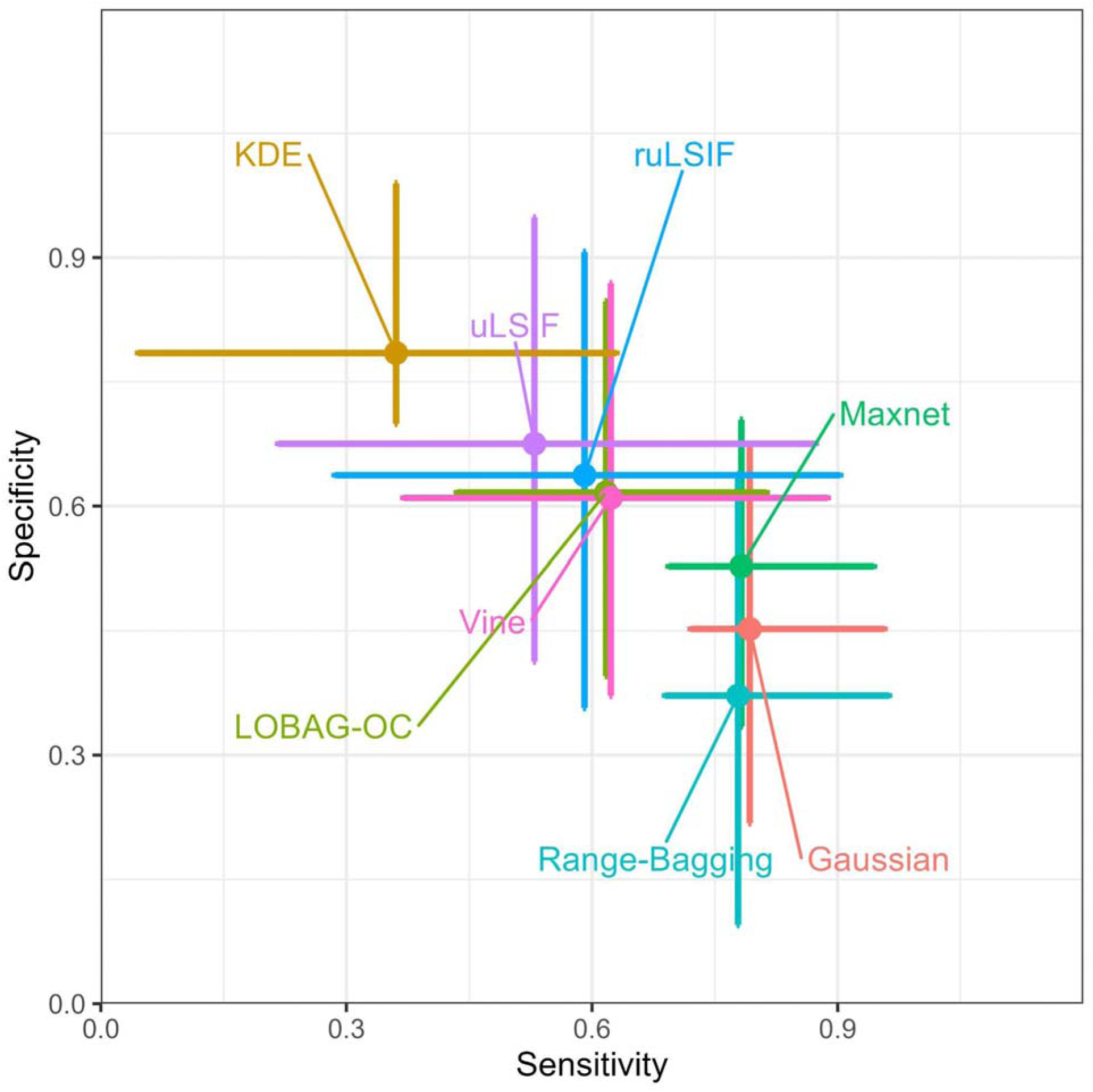
Models vary across a sensitivity-specificity gradient. Variation in model sensitivity and specificity captures variation in assumptions about niche shape and sampling biases. By choosing models that vary along the sensitivity-specificity gradient, we can represent uncertainty in range estimates. Plug-and-play data are categorized by their presence method and include all background methods (see Figure SI 4 for details on performance variation due to background methods).. Points represent median model performance, lines represent interquartile ranges. Models were fit on training data and the resulting ROR were converted to binary presence/absence data by selecting a threshold that omitted 5% of the training presences. Sensitivity and specificity were assessed on independent presence-absence data. See Methods for full details.

Here, we evaluate the performance of different plug-and-play, density-ratio, and environmental-range models across multiple species with varying numbers of presence records using a classic, standardized and vetted dataset in the SDM literature which contains both presence-background data and independent presence-absence data (Elith et al., 2006, 2020). For plug-and-play models, we consider both those that use the same algorithms for estimating both presence and background distributions, as well as those using different algorithms for estimating each distribution. We compare model performances to Maxnet (Maxent implemented via glmnet; Friedman et al., 2010; S. Phillips, 2021) which serves as a useful point of comparison because of its common usage and high performance in previous bakeoffs (Elith et al., 2006; Valavi et al., 2022). Although our focus here is on small (*n* = 1-20), potentially biased, samples, we test the performance of models across a range of sample sizes. While there are many algorithms that perform adequately with large, unbiased samples, how best to model small and biased samples remains poorly understood.

## Methods

### Analyses

#### Data Sources

Occurrence data were taken from Elith *et al*. (2020), a dataset consisting of 226 species across 4 taxonomic groups (plants, birds, bats, and reptiles) and six biogeographic regions (Australian wet tropics; New South Wales, Australia; Ontario, Canada; New Zealand; Switzerland; and South America). This dataset is well-suited to our purposes, as 1) it includes both presence-background data and independent presence-absence data; and 2) it has been previously used to evaluate the relative predictive performance of many SDM algorithms (Elith et al., 2006). As some of our algorithms are only suited to continuous predictors, we excluded any categorical predictors.

An additional set of occurrence records for 1,620 poorly to moderately sampled species (2 to 100 occurrence records) occurring in the state of Florida in the United States (n = 1,620) was downloaded from the BIEN database (B. S. Maitner et al., 2017). Additional climate data were taken from the Worldclim dataset (Hijmans et al., 2005).

### Plug-and-play model comparison

To evaluate the performance of different plug-and-play models, we selected a set of five distribution estimators that varied in the sensitivity-specificity tradeoff: range-bagging (Drake, 2015), LOBAG-OC (Drake, 2014), gaussian (Drake & Richards, 2018), vine copula (Ghosh et al., 2019), and KDE (Li & Racine, 2003). We performed a full factorial design with each algorithm being used to estimate both the presence and background distributions (Table SI 2). We also included each of the five distribution algorithms as a presence-only model (*i.e.*, assuming a uniform background density). The two environmental-range estimators (range-bagging and LOBAG-OC) were not designed for use in a plug-and-play scenario and have not been used in this way in the past. However, the outputs of these estimators could be interpreted probabilistically as uniform distributions over an estimated range. Given that our focus was on data-poor species and that the uniform distributions make few assumptions about the data, we decided to include both range-bagging and LOBAG-OC as components of the plug-and-play models as well as in their original, environmental-range context. Finally, we included three density-ratio algorithms: uLSIF, ruLSIF, and Maxnet (Kanamori et al., 2009; S. Phillips, 2021; Yamada et al., 2013).

## Algorithms considered

### Environmental-Range Algorithms

#### Range-bagging

(Drake, 2015) uses nonparametric bootstrapping to estimate the support of a set of numbers, i.e., its *range*. For each replicate, a subset of environmental predictors and occurrence records are chosen. A convex hull is then constructed around the subset of occurrence records, with locations within the convex hull considered within the niche. This procedure is repeated across bootstrap replicates. The full range-bagging model thus consists of a set of convex hulls in environmental space. To evaluate whether a given point in environmental space is within the niche or not, that point is then compared with the full set of convex hulls, with each comparison resulting in a vote. Thus, for a given environmental distribution, range bagging quantifies the fraction of votes for each location being within a species’ niche.

#### Model settings

Range-bagging has three model parameters: the number of votes to use, the number of covariates to use for each replicate, and the fraction of observations to use for each replicate. We set the number of votes to 100 as previous work showed that there was little improvement in model performance beyond this point (Drake, 2015). The number of environmental dimensions was set to two because using a single dimension has been shown to lead to a dependency of model performance on the fraction of records included in each replicate (Drake, 2015). Including more dimensions would restrict the number of species for which modeling could be done. Further, selecting only two dimensions limits the estimators to only first order interactions, which may be more appropriate for data-poor species. The fraction of observations to use in each replicate was set to 0.5, as this has been shown to provide reasonable AUC scores (Drake, 2015), while also allowing for small sample sizes to be used (*i.e.*, *n* >= 5).

### LOBAG-OC

(Drake, 2014), like range-bagging, relies upon a bootstrapping procedure to characterize species’ niches. However, where range-bagging utilizes convex hulls to delimit niches within each replicate, LOBAG-OC utilizes kernelized hyperspheres (sphere-shaped decision boundaries in n-dimensional space; Tax & Duin, 1999). Further, LOBAG-OC utilizes the full set of environmental space for each bootstrap replicate, where range-bagging typically uses a subset.

#### Model settings

LOBAG-OC has three model parameters: the number of votes to use, and tuning parameters ν(which controls the upper bound on the training error and lower bound on the fraction of data points which become support vectors) and σ(which controls the width of the gaussian kernel used in identifying hyperspheres). We set the number of votes to 100, as previous work has shown diminishing returns on AUC beyond this point (Drake, 2014). We set ν=0.01 as this value has been shown to maximize AUC (Drake, 2014). The value of σ was estimated as part of model fitting.

### Density Algorithms

**Gaussian** estimation entails fitting data to a normal distribution. In the context of SDM, this usually entails fitting a multivariate Gaussian model, which is a generalization of the univariate Gaussian to higher dimensions.

#### Model settings

Drake and Richards (2018) tested three alternative approaches for fitting a Gaussian distribution: ordinary Gaussian estimation, robust estimation, and a regularized approach utilizing a shrinkage estimator, finding that the regularized approach resulted in the highest performance in terms of AUC scores (Drake & Richards, 2018). We thus selected the regularized approach for use in our analyses.

### KDE

(Li & Racine, 2003) is a non-parametric method that estimates probability distributions using kernels as weighting functions. We included KDE because it is a non-parametric method that makes no assumptions about the shape of the underlying distribution. Further, this method can accommodate holes in species distributions.

#### Model settings

Following Drake and Richards (2018), we estimate bandwidth using the method of Li and Racine (2003). Drake and Richards found that other bandwidth estimators lead to massive increases in computation time with only minor gains in performance.

**Vine Copula** models represent a multivariate distribution as a combination of univariate distributions plus copulas that capture the dependencies between variables (Ghosh et al., 2019). Vine copulas were included here due to their ability to handle multivariate distributions with complex dependencies (Ghosh et al., 2019).

#### Model settings

Vine Copula model parameters followed the default recommendations for the vine function in the R package rvinecopulib *(Nagler & Vatter, 2023)*.

### Density Ratio Algorithms

**uLSIF** (Kanamori et al., 2009) is a method that can directly estimate the ratio of two density functions without estimating the component density functions. It does this by recasting the optimization as a least-squares estimation, allowing for solving using quadratic program solvers. This method has been previously found to have lower AUC scores than Maxent (Drake & Richards, 2018), but is included here to provide a point of comparison with a new extension of uLSIF, ruLSIF.

#### Model settings

uLSIF model settings followed the default recommendations for the uLSIF function in the R package densratio (Makiyama, 2019).

**ruLSIF** (Yamada et al., 2013) is an extension of uLSIF that is intended to improve model fitting speed and performance by estimating relative density ratios, which are smoother than ordinary density ratios. This algorithm was included as it has not yet been evaluated in a SDM context.

#### Model settings

uLSIF model settings followed the default recommendations for the RuLSIF function in the R package *densratio* (Makiyama, 2019)

**Maxnet** (S. Phillips, 2021) is an implementation of the Maxent algorithm (S. J. Phillips et al., 2006) that uses glmnet (Friedman et al., 2010) for model fitting. Maxnet serves as a useful point of comparison, as it is among the most popular and best-perming SDM methods (Elith et al., 2006; Valavi et al., 2022).

#### Model settings

Maxnet models were fit with default settings (S. Phillips, 2021).

### Model fitting and evaluation

Plug-and-play models were fitted using the functions fit_plug_and_play() and fit_density_ratio() in the S4DM R package (B. S. Maitner et al., 2025). Continuous model predictions were converted to binary predictions by thresholding the continuous ROR rasters based on the 5% quantile of predicted values at training presences.

The performance of each model was evaluated using 1) the best-case evaluation scenario, an independent presence-absence dataset; 2) 5-fold cross-validation of the training dataset to provide a measure of performance in practical scenario where independent data are not available but there are sufficient data for cross-validation; and 3) the full training dataset as this may be the only option available for especially small sample sizes where subsetting is not feasible. The full models were evaluated using the area under the receiver operating curve (AUC) and correlation between binary presence-background (or presence-absence) and relative occurrence rate (COR; because these are the most common metrics used), and partial AUCs (pAUC; sensitivity ranging from 0.8 to 1; because we rarely care about models with sensitivity < 0.8 in SDM, so this provides a more informative performance measure than AUC). The cross-validated models and full models were evaluated using training AUC and pAUC, test AUC and pAUC, sensitivity, specificity, diagnostic odds ratio (DOR), prediction accuracy, and kappa. We chose to include a diversity of metrics because they capture different elements of model performance (Fielding & Bell, 1997; Liu et al., 2011), and to facilitate comparisons with other work (*e.g.*, Drake, 2015; Drake & Richards, 2018; Elith et al., 2006; Valavi et al., 2022). The presence-absence AUC, COR, and Kappa were also calculated by Elith et al. (2006), allowing comparison of our methods to theirs, however, we note that their Kappa statistic is calculated as the maximum possible Kappa where ours was calculated based the the threshold (five percent training presence omission) that we used. To identify models performing significantly different from Maxnet, we compared the distributions of model AUCs using Mann-Whitney tests (Mann & Whitney, 1947; R Core Team, 2023).

### Small Sample Sizes and Model Performance

As model performance is expected to increase with sample size and we were especially interested in modeling species with small sample sizes, we conducted a second comparison for the subset of species with 20 or fewer occurrence records. For this subset of data-deficient species, we again tested which models performed similarly to Maxnet using Mann-Whitney tests.

For a given species and algorithm, models are also expected to vary in how generalizable they are to different datasets, with generalizability depending on the metrics used to quantify model performance. Even though a model performs well for a particular dataset, it may perform poorly when used in a new region or with a new set of data. For sufficiently large data, cross-validation can be used to quantify model transferability, aiding in model selection (Roberts et al., 2017; Wenger & Olden, 2012). However, cross-validation might be misleading for very small sample sizes. Thus, for data-deficient species, one needs performance metrics which can be reliably estimated from a small sample. To quantify the generality of performance metrics for species with small sample sizes, we examined correlations between model performance metrics quantified using training data vs. those quantified using independent presence-absence data.

For these assessments, we limited our analyses to 34 species with 20 or fewer occurrence records.

### Ensembles to deal with uncertainty

To demonstrate the ability of ensembles to cope with small sample sizes, we evaluated the performance of an ensemble of three algorithms that we found to perform well across sample sizes and which spanned the sensitivity-specificity gradient (Figure 2): Maxnet, KDE/KDE, and ruLSIF. Further, by choosing a subset of algorithms we hoped to balance among their strengths. For each species in the Elith et al. (2020) dataset with 20 or fewer occurrences, we created an ensemble of these three models. There are two ways in which ensembles can be helpful when modeling species. First, they may perform better than individual models (Hao et al., 2020; Thuiller et al., 2009; Valavi et al., 2022). Second, selecting models that vary in their assumptions allows us to capture variation across different model predictions (Buisson et al., 2010; Diniz-Filho et al., 2009). We thus took two approaches to ensembling: model averaging (which ignores variability across models) and vote counting (which preserves variation across models). For the model averaging approach, estimated relative occurrence rates were standardized to sum to one over all locations, and an unweighted average was calculated. We did not weight by AUC (*e.g.*, as in Hao et al., 2020), as data-poor species may not have enough records for accurate estimation of AUC. These average ROR predictions were then thresholded at the 5% quantile of predicted values at presence locations to derive binary labels. For the vote counting approach, individual model predictions were thresholded at the 5% training quantile to yield binary presence/absence predictions. These presence/absence predictions were then summed to produce an ensemble prediction that quantified support for each location in terms of “votes” (*i.e.*, number of models that predicted presence), ranging between zero (full agreement on absence) and three (full agreement on presence).

When evaluating ensemble performance, we selected metrics focused on binary presence/absence classification (*e.g.*, sensitivity, specificity, prediction accuracy), as these can be applied to both the vote counting ensemble and the model averaging ensemble. We compared the performance of ensemble models in three ways to compare performance across different use-cases: 1) presences from the averaged model (which may perform better than individual models), 2) locations with any support in the vote counting ensemble (which may be important if we want to capture any potentially suitable locations), and 3) locations with unanimous support in the vote counting ensemble (which may be important if we want to select locations which are very strongly supported as suitable habitat).

As an example of how ensemble agreement may change with sample size, we created model ensembles for the set of 1,620 plant species with between 2 and 100 occurrence records from BIEN (B. S. Maitner et al., 2017). We used this dataset because it provided many more small sample size species and because assessing model agreement does not require presence-absence data. Predictor variables used were mean annual temperature and mean annual precipitation at a 0.5 degree resolution from the Worldclim dataset (Hijmans et al., 2005). Background data were those within 100 km of an occurrence record.

### R packages used

All analyses were carried out in R (R Core Team, 2023). The S4DM R package (B. S. Maitner et al., 2025) builds upon code from Drake and Richards (2014, 2015; 2018). The S4DM package relies upon the R packages corpcor (Schafer et al., 2021), densratio (Makiyama, 2019), flexclust (Leisch, 2006), geometry (Habel et al., 2023), kernlab (Karatzoglou et al., 2004), maxnet (S. Phillips, 2021), mvtnorm (Genz & Bretz, 2009), np (Hayfield & Racine, 2008), pROC (Robin et al., 2011), robust (Wang et al., 2024), rvinecopulib (Nagler & Vatter, 2023), sf (Pebesma, 2018), terra (Hijmans, 2022), and dplyr (Wickham et al., 2022) and uses the packages geodata (Hijmans et al., 2024), BIEN (B. S. Maitner et al., 2017), ggplot2 (Wickham, 2016), tidyterra (Hernangómez, 2023), knitr (Xie, 2018), and devtools (Wickham et al., 2020) in examples. In our analyses, data are taken from the R packages disdat (Elith et al., 2020), maps (Becker et al., 2023), and geodata (Hijmans et al., 2024), with data wrangling utilizing tidyverse (Wickham et al., 2019). Model fitting and evaluation utilized the packages AUC (Ballings & Van den Poel, 2022), lme4 (Bates et al., 2014), betareg (Cribari-Neto & Zeileis, 2010), bbmle (Bolker & R Development Core Team, 2022), glmmTMB (Brooks et al., 2017), lmerTest (Kuznetsova et al., 2017), nlme (Pinheiro et al., 2020), confintr (Mayer, 2023), DescTools (Signorell, 2024), emmeans (Lenth, 2024), multcomp (Hothorn et al., 2008), margins (Leeper, 2024), and pscl (Zeileis et al., 2008). Parallelization relied upon foreach (Microsoft & Weston, 2022) and doParallel (Microsoft Corporation & Weston, 2022). Plotting and visualization relied upon ggpubr (Kassambara, 2022), tidyterra (Hernangómez, 2023), corrplot (Wei & Simko, 2021), ggpmisc (Aphalo, 2022), lemon (Edwards, 2024), ggnewscale (Campitelli, 2024), plotrix (J, Lemon, 2006), and grid (R Core Team, 2023).

## Results

### Comparative performance: all sample sizes

Across all sample sizes we found that Maxnet models had the highest mean (0.731) and median (0.725) AUCs relative to the independent presence-absence data (Table 1). These AUC scores are comparable to previous studies (Elith et al., 2006). While Maxnet had the highest mean or median AUCs, it only had the highest AUCs for 28% of species, with range-bagging placing second with 7%, gaussian/KDE with 5%, and 21 other models performing best for at least one species (note that throughout we use the convention of presence method / background method; Table SI1). Range-bagging/vine models had the highest sensitivity (0.932), but very poor mean AUC scores (0.507). Presence-only KDE models (*i.e.*, KDE / none) had the highest specificity (0.916) and prediction accuracy (0.819) on the presence-absence data. Kappa relative to the presence-absence dataset was the highest for the KDE/KDE models (0.114). On average, all model types had AUCs that performed better than the random expectation of 0.5. Across all model types, there were three that had AUC distributions not significantly different than Maxnet when assessed for all sample sizes: KDE/KDE (W = 23413.0, p = 0.17; Table SI2), ruLSIF (W = 23198.0, p = 0.11), and Gaussian/KDE (W = 23186.0, p = 0.12). Importantly, even though these three models had AUC values that were similar to Maxnet, sensitivity and specificity differed, suggesting that they achieve a high performance by capturing different information about the distribution (*e.g.*, presences vs absences; Table 1). Median model runtimes varied substantially and ranged from 1s to 5 min (Table SI3).

**Table 1.**
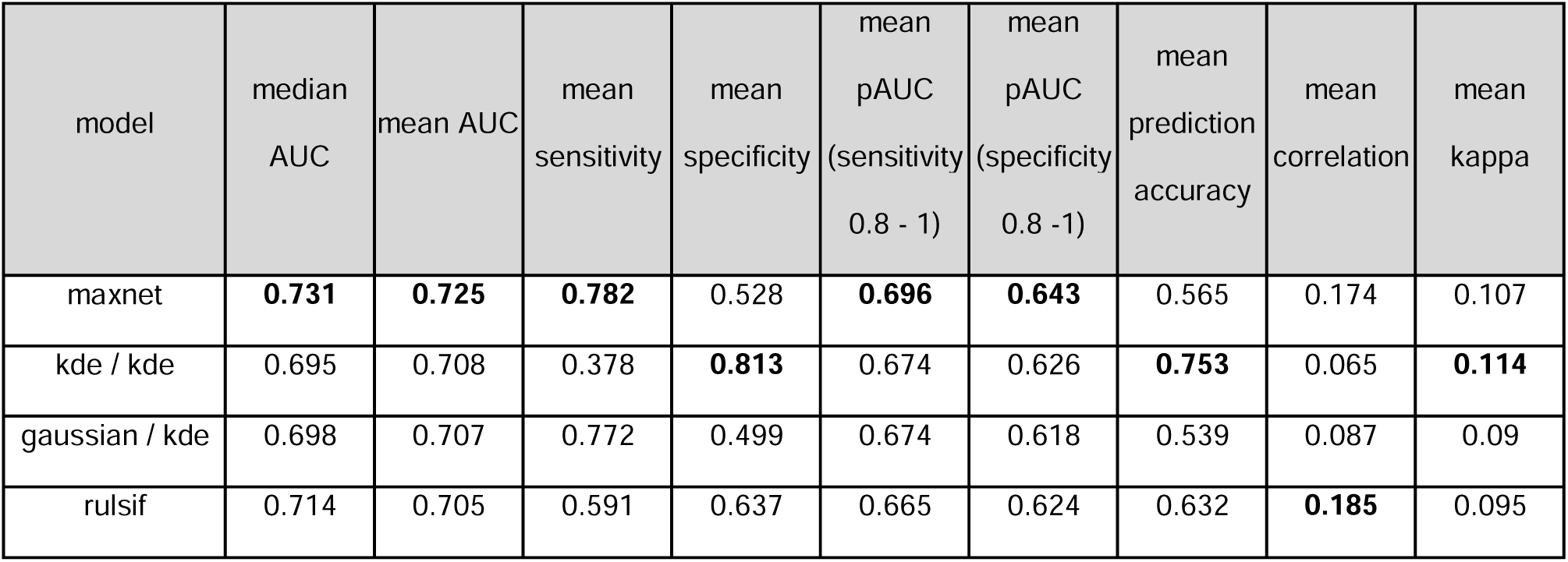
Overall plug-and-play model comparison assessed with presence-absence data. Values shown are the mean values across the entire dataset. Models shown are the subset with AUC distributions not significantly worse than MaxNet (See Table SI1 for the complete set of models). AUC refers to the AUC of the full model, where pAUC refers to the partial AUC. Values highlighted in bold are the best performing models for each performance metric, relative to the set of models with comparable AUC distributions.

### Presence-background model combination performance

Across the different presence-background model combinations, model performance was primarily driven by the presence algorithm used (Figure SI4), with a few key exceptions. In terms of AUC, range-bagging’s performance was best when used either as a presence-only algorithm (assuming uniform background) or when using range-bagging or LOBAG-OC as a background algorithm. In terms of sensitivity and specificity, model performance varied as a function of the background algorithm used when selecting either KDE or range-bagging as a presence algorithm (Figure SI4). When using KDE as a presence algorithm, sensitivity was highest when using a vine algorithm as the background and specificity was high as either a uniform-background algorithm or when using either LOBAG-OC or range-bagging as a background algorithm (Figure SI4).

### Sample size and model performance

#### Relative performance among models: small sample size species

When restricting model comparisons to small sample size species (species with 20 or fewer presence records, *n* = 34 species), the best-performing models for each metric remained largely unchanged (Table 2, Table SI5) with the exception of median AUC (range-bagging) and kappa (LOBAG-OC/LOBAG-OC). Again, all model types had AUCs that performed better than the random expectation of 0.5 on average (Table SI5). Maxnet had the highest presence-absence AUC score for 27% of species, with ruLSIF placing second with 12% and 14 other algorithms winning for at least one of the 34 small sample size species. When focusing only on data-deficient species, the majority of models (24/31) had AUC distributions that did not differ significantly from Maxnet (Table SI6). Notably, all of the models with AUC distributions significantly different from Maxnet’s involved either range-bagging or LOBAG-OC outside of a presence-only context (Table SI6). When using range-bagging as a background function, it performs similarly to assuming a uniform background in most cases (with the notable exceptions of range-bagging / range-bagging; Table 2, SI5). For example, vine / none and vine / rangebagging model performances were similar across all metrics (Table 2, SI5). However, range-bagging is focused on estimating niche edges and is a relatively uniform distribution. When dividing range-bagging presence distributions by a more heterogeneous background distribution (*e.g.*, KDE or Gaussian), the resulting relative occurrence rate is the inverse of the background distribution where the range-bagging distribution takes a value of one and zero where rangebagging takes a value of zero. The predictions of these models thus often assume relatively low occurrence rates in the center of the environmental niche (Figure SI7).

**Table 2.** Model performance evaluated with Presence-Absence data, limited to species 20 or fewer occurrence records. Values shown are the mean values across the entire Elith *et al*. (2020) dataset. Models shown are the subset with AUC distributions not significantly worse than MaxNet (See Table SI2 for the complete set of models). AUC refers to the AUC of the full model, where pAUC refers to the partial AUC. Values highlighted in bold are the best performing models for each performance metric, relative to the set of models with comparable AUC distributions.

| model | median | mean | mean | mean | mean pAUC | mean pAUC | mean | mean | mean |
| --- | --- | --- | --- | --- | --- | --- | --- | --- | --- |

|  | AUC | AUC | sensitivity | specificity | (sensitivity<br>0.8 - 1) | (specificity 0.8<br>- 1) | prediction<br>accuracy | correlation | kappa |
| --- | --- | --- | --- | --- | --- | --- | --- | --- | --- |
| maxnet | 0.656 | **0.697** | **0.593** | 0.667 | **0.712** | **0.67** | 0.651 | 0.163 | 0.11 |
| rangebagging /<br>rangebagging | **0.68** | 0.692 | 0.164 | 0.949 | 0.645 | 0.629 | 0.833 | **0.177** | 0.075 |
| vine / gaussian | 0.635 | 0.682 | 0.231 | 0.902 | 0.644 | 0.638 | 0.799 | 0.064 | 0.105 |
| vine / kde | 0.631 | 0.68 | 0.213 | 0.903 | 0.644 | 0.619 | 0.8 | 0.094 | 0.099 |
| gaussian / kde | 0.662 | 0.678 | 0.473 | 0.722 | 0.653 | 0.61 | 0.682 | 0.055 | 0.093 |
| vine / vine | 0.634 | 0.677 | 0.285 | 0.854 | 0.643 | 0.625 | 0.79 | 0.084 | **0.111** |
| kde / kde | 0.658 | 0.677 | 0.089 | 0.97 | 0.667 | 0.625 | 0.82 | 0.09 | 0.061 |
| vine / none | 0.639 | 0.676 | 0.213 | 0.899 | 0.641 | 0.619 | 0.796 | 0.099 | 0.092 |
| vine /<br>rangebagging | 0.639 | 0.676 | 0.213 | 0.899 | 0.641 | 0.619 | 0.796 | 0.099 | 0.092 |
| rulsif | 0.679 | 0.675 | 0.348 | 0.791 | 0.649 | 0.618 | 0.725 | 0.151 | 0.081 |
| kde / none | 0.656 | 0.674 | 0.012 | **0.998** | 0.655 | 0.623 | **0.837** | 0.094 | 0.013 |
| kde /<br>rangebagging | 0.656 | 0.674 | 0.012 | **0.998** | 0.655 | 0.623 | **0.837** | 0.094 | 0.013 |
| rangebagging /<br>none | 0.666 | 0.673 | 0.177 | 0.943 | 0.645 | 0.627 | 0.821 | 0.163 | 0.076 |
| kde / gaussian | 0.659 | 0.669 | 0.172 | 0.906 | 0.664 | 0.638 | 0.782 | 0.048 | 0.065 |
| ulsif | 0.658 | 0.668 | 0.277 | 0.865 | 0.636 | 0.622 | 0.784 | 0.145 | 0.08 |
| gaussian / none | 0.655 | 0.667 | 0.477 | 0.705 | 0.633 | 0.613 | 0.669 | 0.125 | 0.09 |
| gaussian /<br>rangebagging | 0.655 | 0.667 | 0.474 | 0.706 | 0.633 | 0.613 | 0.669 | 0.125 | 0.09 |
| gaussian /<br>gaussian | 0.624 | 0.662 | 0.516 | 0.692 | 0.682 | 0.625 | 0.67 | 0.024 | 0.089 |
| kde / vine | 0.616 | 0.659 | 0.221 | 0.842 | 0.657 | 0.624 | 0.763 | 0.032 | 0.064 |
| vine / lobagoc | 0.65 | 0.653 | 0.245 | 0.825 | 0.611 | 0.594 | 0.724 | 0.067 | 0.07 |
| kde / lobagoc | 0.619 | 0.648 | 0.103 | 0.91 | 0.626 | 0.615 | 0.751 | 0.092 | 0.017 |
| rangebagging / | 0.626 | 0.646 | 0.258 | 0.822 | 0.634 | 0.619 | 0.712 | 0.127 | 0.049 |
| **lobagoc** |  |  |  |  |  |  |  |  |  |
| **gaussian / vine** | 0.596 | 0.646 | 0.575 | 0.606 | 0.645 | 0.606 | 0.631 | 0.02 | 0.077 |
| **gaussian /**<br>**lobagoc** | 0.642 | 0.646 | 0.5 | 0.645 | 0.616 | 0.61 | 0.609 | 0.124 | 0.085 |
| **lobagoc / none** | 0.595 | 0.606 | 0.266 | 0.868 | 0.542 | 0.597 | 0.786 | 0.144 | 0.108 |

#### Assessing model quality predictability

One of the challenges with assessing model performance for poorly-sampled species is that there may be insufficient records to allow robust cross-validation of model performance. We thus tested for correlations between performance calculated relative to training data and performance relative to independent, presence-absence data. As a metric of overall model reliability, we also calculated the percentage of models which resulted in AUC values worse than random (*i.e*, AUC < 0.5). Specificity and prediction accuracy showed strong correlations between values obtained on training data and those obtained on independent, presence-absence data (*r* = 0.91, p < 2.2 e-16 and *r* = 0.77, p < 2.2 e-16, respectively; Figure SI8). AUC, sensitivity, and COR showed significant but weak correlations (*r* = 0.22, p < 8 e-13; *r* = 0.11, p = 0.00024; *r* = 0.067, p = 0.03; respectively) while kappa was significant, weak, and negative (*r* = -0.074, p = 0.016). Maxnet produced models with AUC values worse than random in 15% of cases, while 19 algorithms performed as well or better than Maxnet, including several using Gaussian distributions, KDE, vine copulas, and range-bagging as presence methods (Table SI9). In cases where there were enough points for cross-validation of poorly-sampled species (n = 32 species), we observed only weak to moderate correlation between mean testing AUC values and AUC values observed in the independent, presence-absence data for both AUC score (r = 0.36, p = 2.2e-16) and relative AUC ranking (ρ = 0.23, p = 1.542e-13). Only in 16% of cases was the best performing model in terms of test AUC the same model in terms of independent, presence-absence AUC. This suggests that, for species with small sample sizes, models performing well in terms of cross-validated AUC don’t necessarily provide a reliable estimate of a species’ true range, as they might be capturing patterns specific to the training dataset rather than generalizable ecological relationships. Further, relying solely on the model that tends to perform best overall in terms of mean AUC (Maxnet) may be unwise, as it may result in models that are worse than random, and is rarely the best performing model on any given species.

### Model Ensembles

Due to the challenges associated with identifying a single best model overall, we explored ensembles as a way to improve performance and capture model disagreement. For our ensembles, we selected three models with AUC distributions comparable to Maxnet at all sample sizes, but spanning the full range of the sensitivity-specificity gradient: KDE/KDE (high specificity, low sensitivity), ruLSIF (intermediate specificity and sensitivity), and Maxnet (low specificity, high sensitivity). Ensembling via a vote counting approach yielded improved performance over component models in terms of sensitivity, specificity, and prediction accuracy. Where presences were taken as locations with support from all models (Ensemble unanimous support, Figure 3, Table SI10), specificity and prediction accuracy were higher than for any of the component models. Where presences were taken as locations with support from any models, sensitivity was higher than for any of the component models (Ensemble any support, Figure 3, Table SI10). However, Maxnet had the highest mean kappa (Table SI10).

**Figure 3.**
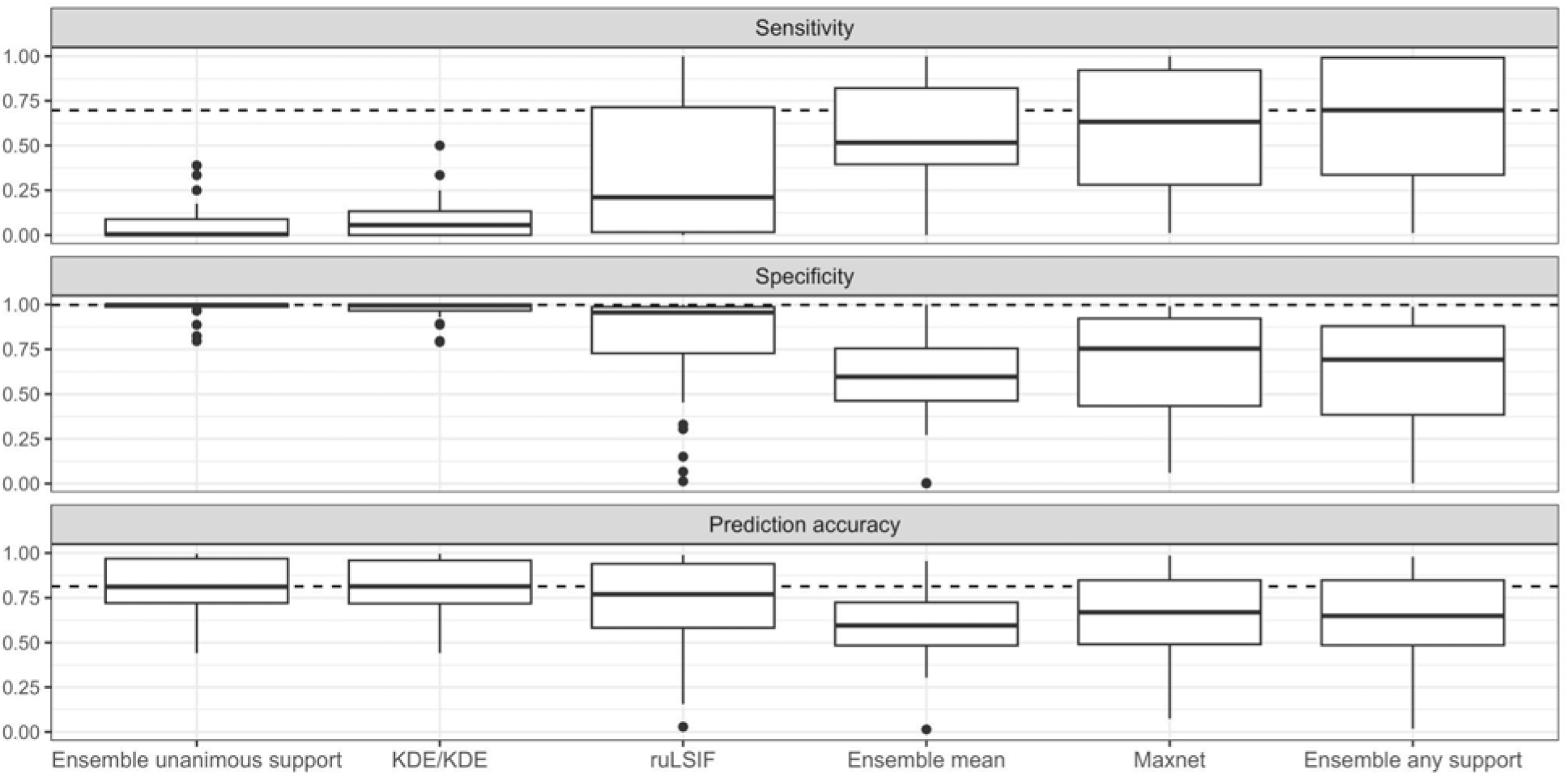
Ensemble performance relative to component models. Ensembles were composed of KDE/KDE, ruLSIF, and Maxnet algorithms. Ensembling was done in two ways: 1) by averaging of model predictions, followed by thresholding (resulting in binary predictions); and 2) by thresholding of individual models followed by aggregation (resulting in support for each location between 0 and 3 votes). The “ensemble unanimous support” and “ensemble any support” refer to considering locations which all models agree on as presence or any models agree on as presence, respectively. Only species with 20 or fewer occurrence records were included.

Agreement across models within an ensemble increased as a function of the number of occurrence records available (Figure 4). When only two occurrence records were available, the three component models agreed for only 18% of predicted locations on average, with 52% of locations receiving support from only one model. However, when 100 occurrence records were available, the models agreed for 69% of locations on average with only 9% of locations receiving support from only one model. Model consensus increased most rapidly between 2 and 25 occurrence records, beyond which point the models tended to agree on most (>50%) predicted locations (Figure 4). We note that our relatively low model agreement reflects our choice of models that span the sensitivity-specificity gradient, and which will thus tend to disagree.

**Figure 4.**
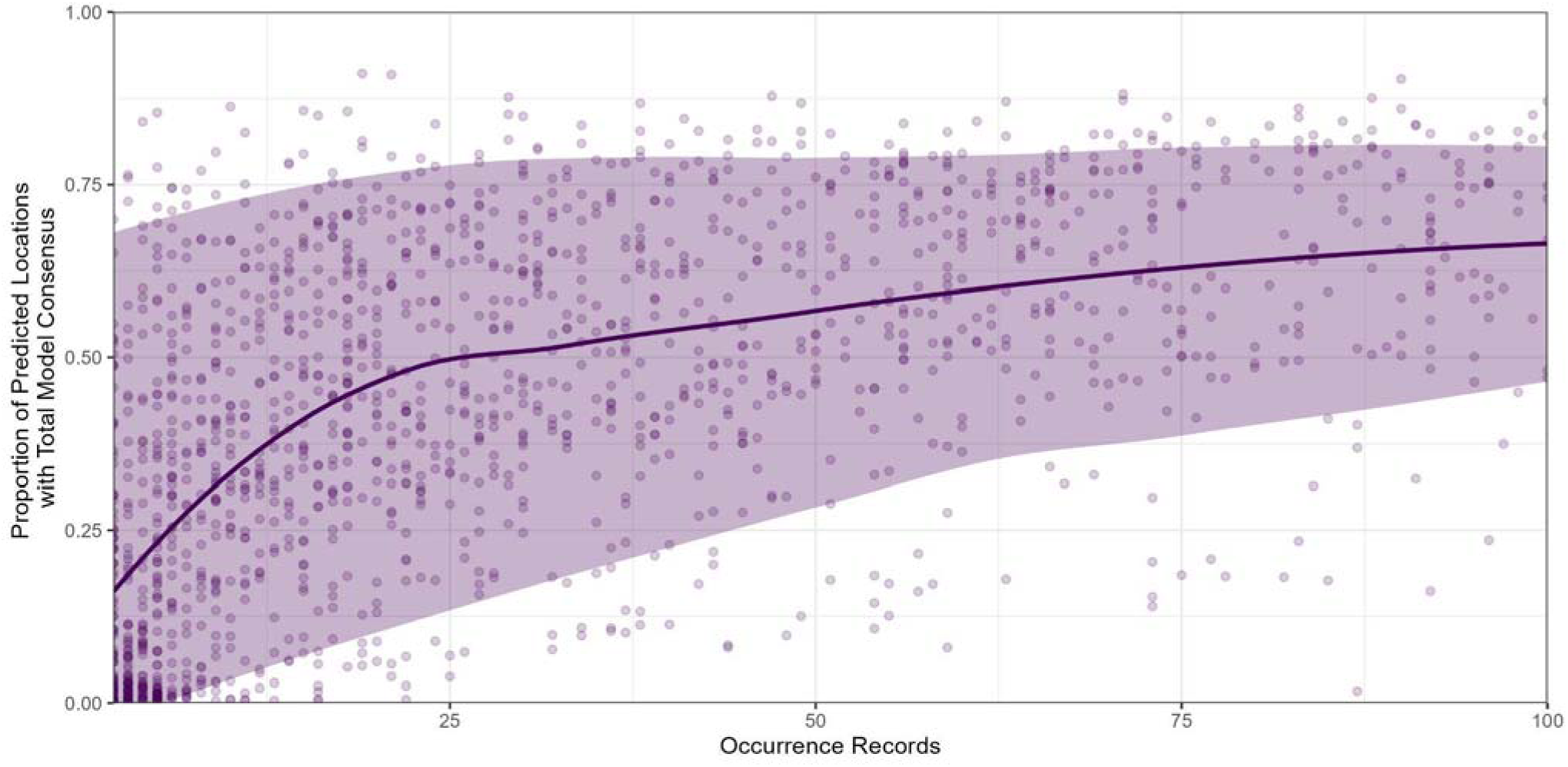
Ensemble agreement increases with sample size. Ensemble agreement was calculated for each species as the number of grid cells with support from all models divided by the number of grid cells with support from at least one model. Ensembles were calculated with KDE/KDE, ruLSIF, and Maxnet algorithms for 1,620 occurrence records of plant species in Florida obtained from BIEN (B. S. Maitner et al., 2017). The solid line is a loess fit to the mean, while the shaded region is the 95% confidence interval defined by loess fits to the 0.025 and 0.975 quantiles. For simplicity, models were calculated using only two variables, mean annual temperature and mean annual precipitation from BioClim (Hijmans et al., 2005). .

## Discussion

When looking across all species (*n* = 226), Maxnet tended to perform the best in terms of AUC, however the algorithms KDE/KDE, Gaussian/KDE, and ruLSIF had similar AUC distributions.

While Maxnet tended to perform the best on average across all species, it only performed the best for 28% of species, with one or more of the alternative methods outperforming Maxnet for 72% of species, suggesting that these alternative methods are useful to consider.

When considering only the small sample size species (*n* = 34), Maxnet performed best in terms of overall AUC, but again, only for 27% of species. In most cases (73%), one or more of the alternatives outperformed Maxnet. Further, the majority of algorithms (77%) had AUC distributions that were not significantly different from Maxnet, supporting the idea that these alternative algorithms may be just as useful as Maxnet for small sample size species. While these models had similar AUC scores, they differed widely in their predictions, which spanned a gradient of sensitivity and specificity. Our results indicate that it will be difficult to select the best performing model from among these viable alternative methods. For some cases, data-poor species will have too few records for cross validation. For cases where cross-validation is possible, the testing sample sizes will be small, increasing variation in the AUC among folds. Further, our results suggest that most performance metrics (aside from specificity and prediction accuracy) estimated from training data for data-poor species are not strong predictors of model predictive performance. Correlations between testing AUC and independent, presence-absence AUC were weak to moderate, suggesting that even when cross-validation is possible for data-deficient species, it may be of limited utility in distinguishing among models. Thus, in the absence of additional knowledge of sampling assumptions or particular use-cases that might inform model selection, identifying the best model for poorly-sampled species is difficult.

To address this difficulty, we investigated ensembles as a way of accounting for uncertainty in model performance and sampling assumptions. We considered an ensemble of three models that spanned the sensitivity-specificity gradient, and compared vote-counting and model-averaging approaches to ensembling. We found that the performance (in terms of sensitivity, specificity, and prediction accuracy) of the vote-counting ensembles were the highest, and outperformed the component models. Specificity was maximized when selecting locations that were included by all models, while sensitivity was maximized by selecting locations that were included by any models. In the cases where samples do a good job of representing the distribution, the high specificity approach will be more appropriate. In cases where samples are a subset of the distribution, the high sensitivity approach will be more appropriate. In some cases there will be additional information available (*e.g.*, expert opinion on sampling completeness) to help choose between high-sensitivity and high-specificity approaches. However, in most cases the sampling intensity is not known (or, in some cases, not even well-defined, as in compilations of records from heterogeneous sources or citizen science data). By considering a range of possibilities via an ensemble, we can capture this underlying uncertainty using a vote-counting approach that provides upper and lower estimates on species ranges, as well as a measure of relative support for different locations. The importance of incorporating this model disagreement decreases with sample size, as we found that the predictions of different algorithms converged as sample sizes increased (Figure 4).

The modeling frameworks we follow here were first proposed by Drake and Richards (Drake, 2014, 2015; Drake & Richards, 2018). In the initial publications introducing LOBAG-OC and range-bagging algorithms, Drake found their performance to be comparable to Maxent, while later work by Drake and Richards (2018) found that Maxent tended to outperform either range-bagging or LOBAG-OC. Here, we find that, while Maxnet tends to perform better than either LOBAG-OC or range-bagging in terms of AUC evaluated on independent presence-absence data, these algorithms do outperform Maxnet for a subset of species (20% and 35%, respectively). Drake and Richards (2018) also found that KDE/KDE and Gaussian/Gaussian algorithms performed comparably to Maxent, with uLSIF performing poorer. While our results also show a comparable performance between KDE/KDE and Maxent, we find that Gaussian/KDE hybrid models actually tend to perform better than Gaussian/Gaussian models. Our findings supported the poorer performance of uLSIF vs. Maxnet, but ruLSIF, a modification of uLSIF, performed comparably to Maxnet. Where the work of Drake and Richards focused on plug-and-play models that used the same algorithm for both the numerator and denominator, here we additionally explored “hybrid” plug-and-play models which used differing algorithms for the presence and background distribution. We found that these hybrids often outperformed those studied by Drake *et al*. previously (*e.g.*, Gaussian/KDE outperformed Gaussian/Gaussian). This not only confirms the utility of these hybrid models, but suggests that understanding when to use different component algorithms may be a fruitful avenue for improving SDMs generally, and in particular for poorly-sampled species. Previous studies on SDMs have found that ensembles often improve model predictions (Hao et al., 2020; Thuiller et al., 2009; Valavi et al., 2022). However, this improved performance may require model tuning (Valavi et al., 2022) which may not be feasible for poorly-sampled species, or may be implemented through model averaging, which destroys possibly valuable information about model disagreement. As an alternative, we adopted a vote-counting approach, which preserves model disagreement in the output raster and captures different sampling assumptions and use-cases.

### Recommendations

Although the models presented here are viable options for modeling data-deficient species, model performance may be low in many cases and identifying the best-performing model remains challenging. We recommend running an ensemble of models that perform well on average, run reasonably fast, and suit the use-case at hand. Where one is especially interested in getting the presences correct (*e.g.*, for reserve site selection) or where the sampling adequately captures the range of environments a species occupies (*e.g.*, for well-sampled species that have small niches), it makes sense to focus on an ensemble of models towards the high-specificity side of the sensitivity-specificity gradient (Fig. 2), with particular focus on locations where all models agree. Conversely, if one is interested in getting the absences correct (*e.g.*, inferring potential invasive ranges) or where we’ve likely under-sampled the species across its environment (*e.g.*, for poorly-sampled, widely-dispersed species), it would make sense to focus on an ensemble of models on the high-sensitivity side of the gradient (Fig. 4), with focus on locations supported by any model. Often, we will lack *a priori* knowledge of sampling quality, and will be interested in a balance of sensitivity and specificity (*e.g.,* for macroecology or biogeography). In these cases, we recommend selecting an ensemble of algorithms that span the sensitivity-specificity gradient, followed by analyses either across all component models, or at a minimum conducting analyses across (i) location with support by any models and (ii) locations with support by all models as a way of placing upper and lower bounds on distributions.

### Future Directions

Our work suggests several areas of potential future research and development. As new methods for fitting either distributions or the ratios of distributions become available, these can be implemented within the plug-and-play framework (Drake & Richards, 2018). Indeed, we found some of the algorithms that were introduced after Drake and Richard (2018), including vine copulas and ruLSIF, out-perform earlier approaches. Future studies attempting to understand the variation in algorithm performance across species could be a valuable way of selecting models for particular use cases. Finally, we note that newer, more representative datasets would be valuable for continued methodological development and validation.

## Acknowledgements

BSM, BLC, and CM were supported by NASA Grant 80NSSC 22K0883 and CM was supported by NSF BoCP-2225078.

## Author Contributions

BSM, JMD, and CM conceptualized this work. Software and methodologies were developed by JMD, RR, BSM, and CM. Formal analysis and investigation were done by BSM and CM. Visualization was done by BSM and BLC. The original draft was written by BSM, CM and BLC with all authors contributing to review and editing. CM acquired funding.

## Data Availability

All data used in this work have been previously published and are publicly accessible. All code, tables, figures, and derived data products are publicly available via Github (https://github.com/bmaitner/small_sample_size_sdms) and are permanently archived on Zenodo (https://zenodo.org/records/14902720).

## Supplementary Information

**Table SI 1.**
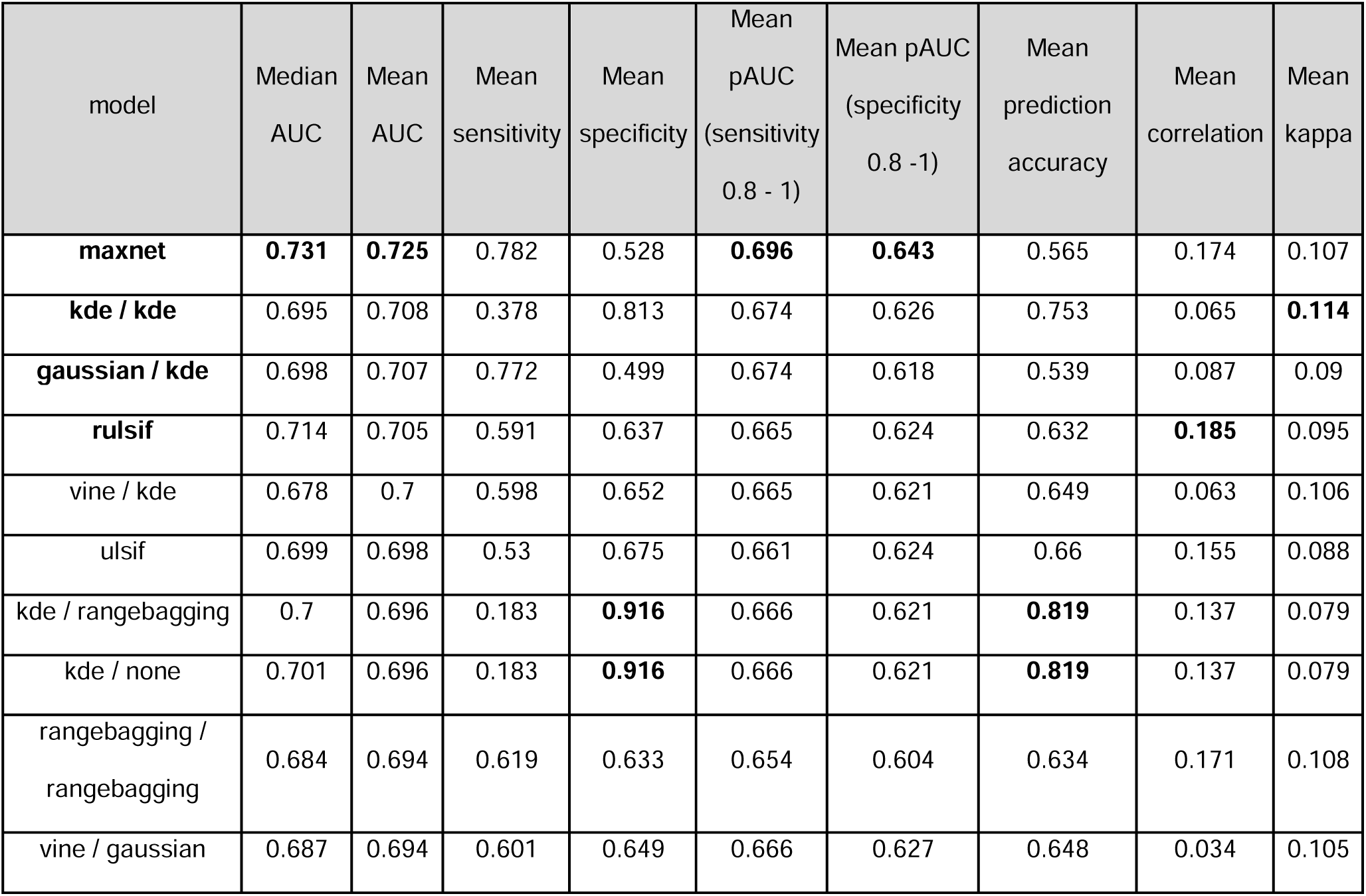
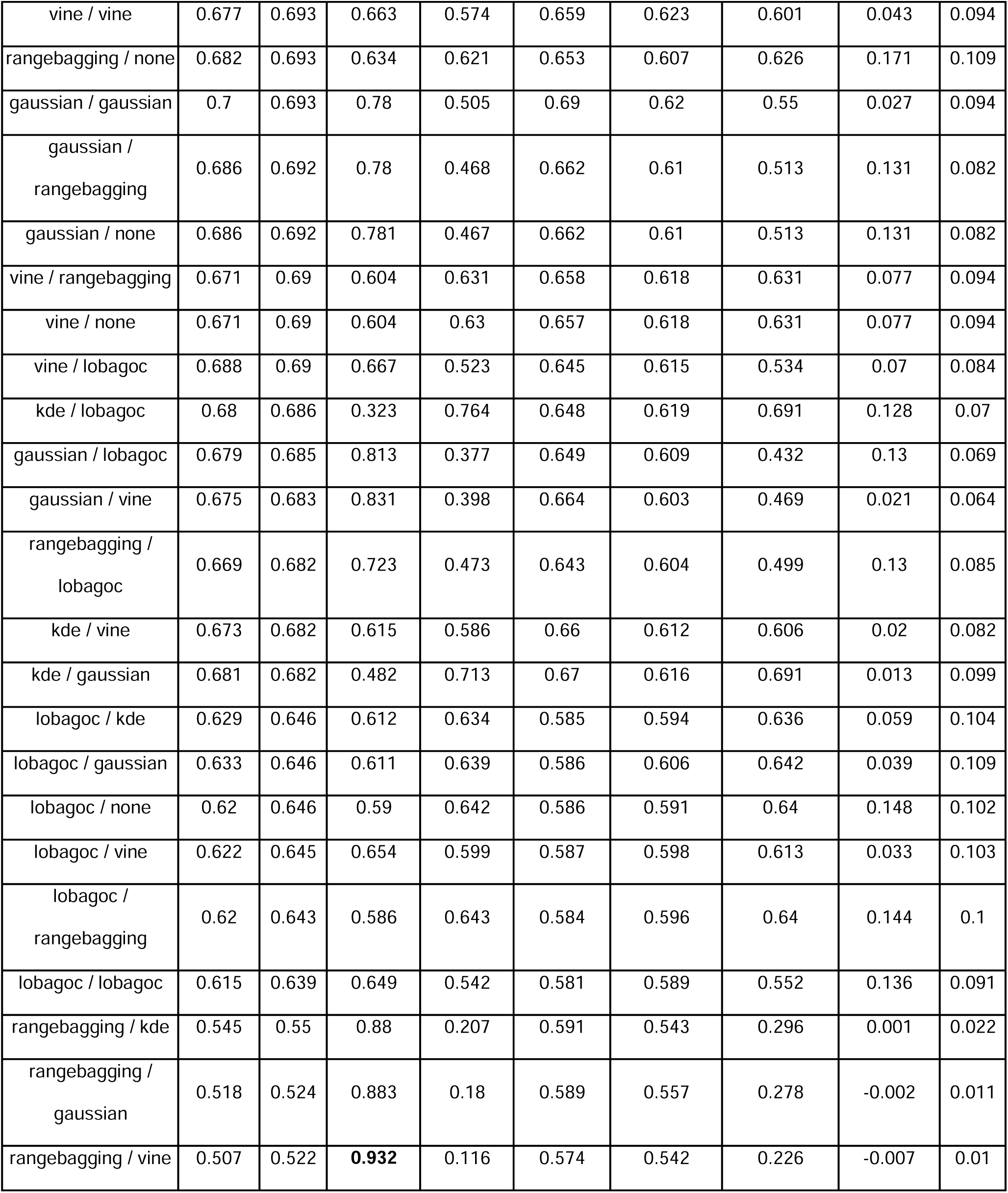
Overall model comparison assessed with presence-absence data. Values shown are the mean values across the entire dataset. AUC refers to the AUC of the full model, where pAUC refers to the partial AUC. Values highlighted in bold are the best performing models for each performance metric. Models highlighted in bold have AUCs not significantly different from Maxnet. See Table 2 for performance limited to SSSS.

| model | Median<br>AUC | Mean<br>AUC | Mean<br>sensitivity | Mean<br>specificity | Mean<br>pAUC<br>(sensitivity<br>0.8 - 1) | Mean pAUC<br>(specificity<br>0.8 - 1) | Mean<br>prediction<br>accuracy | Mean<br>correlation | Mean<br>kappa |
| --- | --- | --- | --- | --- | --- | --- | --- | --- | --- |
| **maxnet** | **0.731** | **0.725** | 0.782 | 0.528 | **0.696** | **0.643** | 0.565 | 0.174 | 0.107 |
| **kde / kde** | 0.695 | 0.708 | 0.378 | 0.813 | 0.674 | 0.626 | 0.753 | 0.065 | **0.114** |
| **gaussian / kde** | 0.698 | 0.707 | 0.772 | 0.499 | 0.674 | 0.618 | 0.539 | 0.087 | 0.09 |
| **rulsif** | 0.714 | 0.705 | 0.591 | 0.637 | 0.665 | 0.624 | 0.632 | **0.185** | 0.095 |
| vine / kde | 0.678 | 0.7 | 0.598 | 0.652 | 0.665 | 0.621 | 0.649 | 0.063 | 0.106 |
| ulsif | 0.699 | 0.698 | 0.53 | 0.675 | 0.661 | 0.624 | 0.66 | 0.155 | 0.088 |
| kde / rangebagging | 0.7 | 0.696 | 0.183 | **0.916** | 0.666 | 0.621 | **0.819** | 0.137 | 0.079 |
| kde / none | 0.701 | 0.696 | 0.183 | **0.916** | 0.666 | 0.621 | **0.819** | 0.137 | 0.079 |
| rangebagging /<br>rangebagging | 0.684 | 0.694 | 0.619 | 0.633 | 0.654 | 0.604 | 0.634 | 0.171 | 0.108 |
| vine / gaussian | 0.687 | 0.694 | 0.601 | 0.649 | 0.666 | 0.627 | 0.648 | 0.034 | 0.105 |
| vine / vine | 0.677 | 0.693 | 0.663 | 0.574 | 0.659 | 0.623 | 0.601 | 0.043 | 0.094 |
| rangebagging / none | 0.682 | 0.693 | 0.634 | 0.621 | 0.653 | 0.607 | 0.626 | 0.171 | 0.109 |
| gaussian / gaussian | 0.7 | 0.693 | 0.78 | 0.505 | 0.69 | 0.62 | 0.55 | 0.027 | 0.094 |
| gaussian /<br>rangebagging | 0.686 | 0.692 | 0.78 | 0.468 | 0.662 | 0.61 | 0.513 | 0.131 | 0.082 |
| gaussian / none | 0.686 | 0.692 | 0.781 | 0.467 | 0.662 | 0.61 | 0.513 | 0.131 | 0.082 |
| vine / rangebagging | 0.671 | 0.69 | 0.604 | 0.631 | 0.658 | 0.618 | 0.631 | 0.077 | 0.094 |
| vine / none | 0.671 | 0.69 | 0.604 | 0.63 | 0.657 | 0.618 | 0.631 | 0.077 | 0.094 |
| vine / lobagoc | 0.688 | 0.69 | 0.667 | 0.523 | 0.645 | 0.615 | 0.534 | 0.07 | 0.084 |
| kde / lobagoc | 0.68 | 0.686 | 0.323 | 0.764 | 0.648 | 0.619 | 0.691 | 0.128 | 0.07 |
| gaussian / lobagoc | 0.679 | 0.685 | 0.813 | 0.377 | 0.649 | 0.609 | 0.432 | 0.13 | 0.069 |
| gaussian / vine | 0.675 | 0.683 | 0.831 | 0.398 | 0.664 | 0.603 | 0.469 | 0.021 | 0.064 |
| rangebagging /<br>lobagoc | 0.669 | 0.682 | 0.723 | 0.473 | 0.643 | 0.604 | 0.499 | 0.13 | 0.085 |
| kde / vine | 0.673 | 0.682 | 0.615 | 0.586 | 0.66 | 0.612 | 0.606 | 0.02 | 0.082 |
| kde / gaussian | 0.681 | 0.682 | 0.482 | 0.713 | 0.67 | 0.616 | 0.691 | 0.013 | 0.099 |
| lobagoc / kde | 0.629 | 0.646 | 0.612 | 0.634 | 0.585 | 0.594 | 0.636 | 0.059 | 0.104 |
| lobagoc / gaussian | 0.633 | 0.646 | 0.611 | 0.639 | 0.586 | 0.606 | 0.642 | 0.039 | 0.109 |
| lobagoc / none | 0.62 | 0.646 | 0.59 | 0.642 | 0.586 | 0.591 | 0.64 | 0.148 | 0.102 |
| lobagoc / vine | 0.622 | 0.645 | 0.654 | 0.599 | 0.587 | 0.598 | 0.613 | 0.033 | 0.103 |
| lobagoc /<br>rangebagging | 0.62 | 0.643 | 0.586 | 0.643 | 0.584 | 0.596 | 0.64 | 0.144 | 0.1 |
| lobagoc / lobagoc | 0.615 | 0.639 | 0.649 | 0.542 | 0.581 | 0.589 | 0.552 | 0.136 | 0.091 |
| rangebagging / kde | 0.545 | 0.55 | 0.88 | 0.207 | 0.591 | 0.543 | 0.296 | 0.001 | 0.022 |
| rangebagging /<br>gaussian | 0.518 | 0.524 | 0.883 | 0.18 | 0.589 | 0.557 | 0.278 | -0.002 | 0.011 |
| rangebagging / vine | 0.507 | 0.522 | **0.932** | 0.116 | 0.574 | 0.542 | 0.226 | -0.007 | 0.01 |

**Table SI 2.** Model comparison with Maxnet. Test statistic (W) and p-value refer to the results of a Mann-Whitney test comparing the AUCs of the focal model with those of Maxnet. Models in bold are NOT significantly different from Maxnet.

| Model | W | p-value |
| --- | --- | --- |
| **maxnet** | **25312.5** | **1** |
| **kde / kde** | **23413** | **0.169** |
| **gaussian / kde** | **23186** | **0.123** |
| **rulsif** | **23198** | **0.108** |
| gaussian / gaussian | 22493 | 0.041 |
| vine / kde | 22281 | 0.041 |
| ulsif | 22544 | 0.037 |
| kde / rangebagging | 22196 | 0.024 |
| kde / none | 22189 | 0.024 |
| vine / gaussian | 21968 | 0.023 |
| vine / vine | 21006 | 0.011 |
| gaussian / rangebagging | 21780 | 0.01 |
| gaussian / none | 21775 | 0.01 |
| rangebagging / rangebagging | 20827 | 0.009 |
| rangebagging / none | 20686 | 0.009 |
| vine / rangebagging | 21366 | 0.007 |
| vine / none | 21337 | 0.006 |
| vine / lobagoc | 21440 | 0.005 |
| kde / gaussian | 21369 | 0.004 |
| kde / lobagoc | 21168.5 | 0.003 |
| gaussian / lobagoc | 21063 | 0.002 |
| gaussian / vine | 19384 | 0.002 |
| kde / vine | 19539 | 0.001 |
| rangebagging / lobagoc | 19842.5 | 0.001 |
| lobagoc / gaussian | 16685 | 0 |
| lobagoc / vine | 16461 | 0 |
| lobagoc / kde | 16698 | 0 |
| lobagoc / none | 16417 | 0 |
| lobagoc / rangebagging | 16397 | 0 |
| lobagoc / lobagoc | 16202 | 0 |
| rangebagging / kde | 8671 | 0 |
| rangebagging / gaussian | 7598 | 0 |
| rangebagging / vine | 6375 | 0 |

**Table SI 3.** Median model runtimes. Model runtime was assessed using the full set of training data. Model fitting time of plug-and-play models is driven primarily by the time needed to fit the background distribution. We note that in cases where multiple species are modelled using a common set of background records, the calculation of the background distribution need only be done once, potentially reducing runtimes by orders of magnitude.

| Method | Runtime (s) |
| --- | --- |
| gaussian / none | 0.002 |
| kde / none | 0.006 |
| gaussian / gaussian | 0.044 |
| kde / gaussian | 0.073 |
| rangebagging / none | 0.119 |
| rangebagging / gaussian | 0.207 |
| lobagoc / none | 0.455 |
| lobagoc / gaussian | 0.465 |
| gaussian / rangebagging | 0.71 |
| kde / rangebagging | 0.805 |
| rangebagging / rangebagging | 0.835 |
| lobagoc / rangebagging | 1.117 |
| vine / gaussian | 1.94 |
| vine / none | 2.016 |
| vine / rangebagging | 2.745 |
| maxnet | 4.966 |
| rulsif | 21.328 |
| lobagoc / lobagoc | 30.98 |
| ulsif | 31.944 |
| rangebagging / lobagoc | 32.309 |
| vine / lobagoc | 34.263 |
| kde / lobagoc | 34.264 |
| gaussian / lobagoc | 37.608 |
| gaussian / kde | 43.359 |
| lobagoc / kde | 44.576 |
| kde / kde | 44.69 |
| rangebagging / kde | 44.943 |
| vine / kde | 45.905 |
| lobagoc / vine | 295.325 |
| gaussian / vine | 300.5 |
| kde / vine | 302.508 |
| vine / vine | 310.251 |
| rangebagging / vine | 310.922 |

**Figure SI 4.**
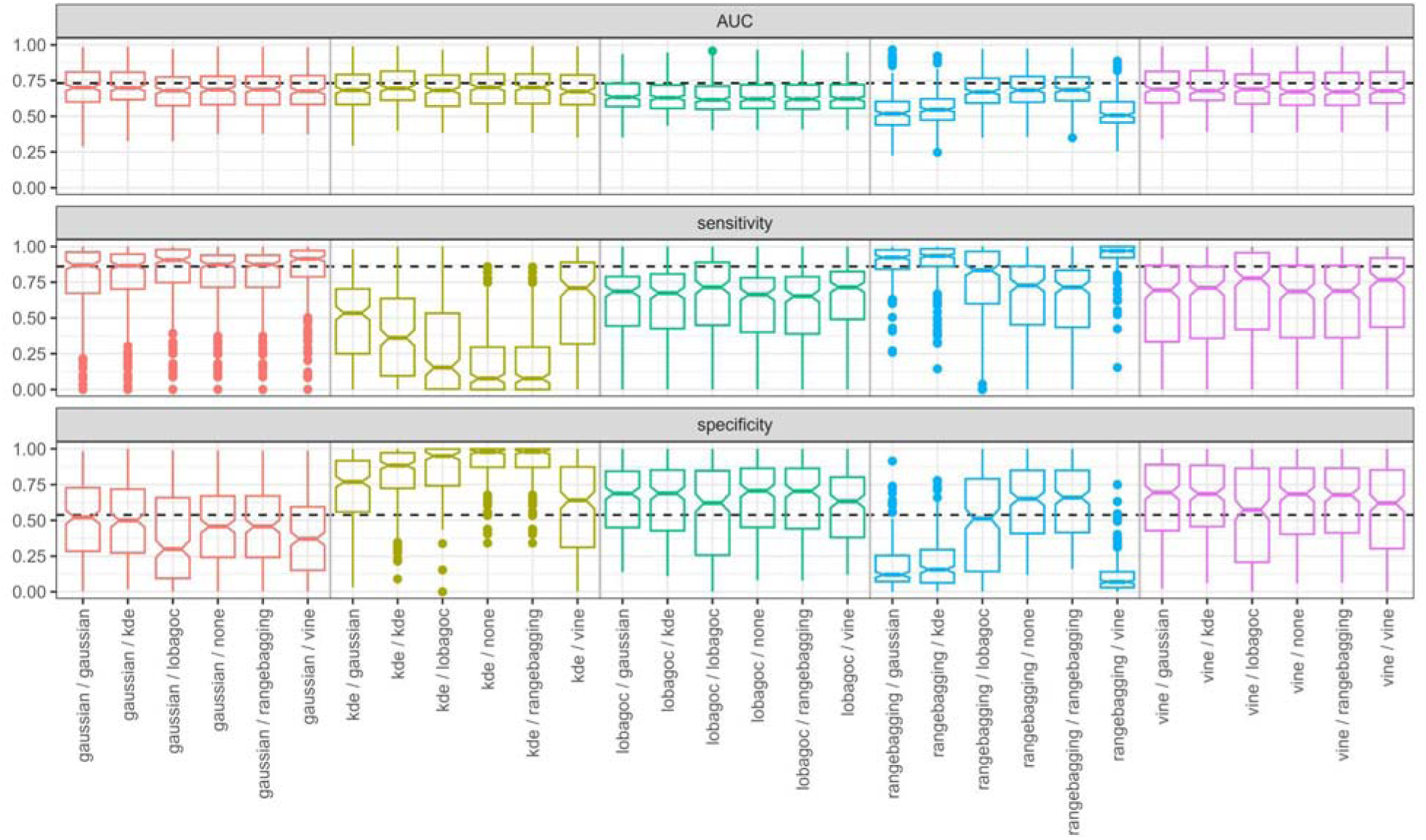
Boxplot of presence-background model performance. AUC, sensitivity, and specificity were calculated relative to independent, presence-absence data. The dashed horizontal lines denote the median values for Maxnet for those metrics. Colors correspond to different presence methods.

**Table SI 5.**
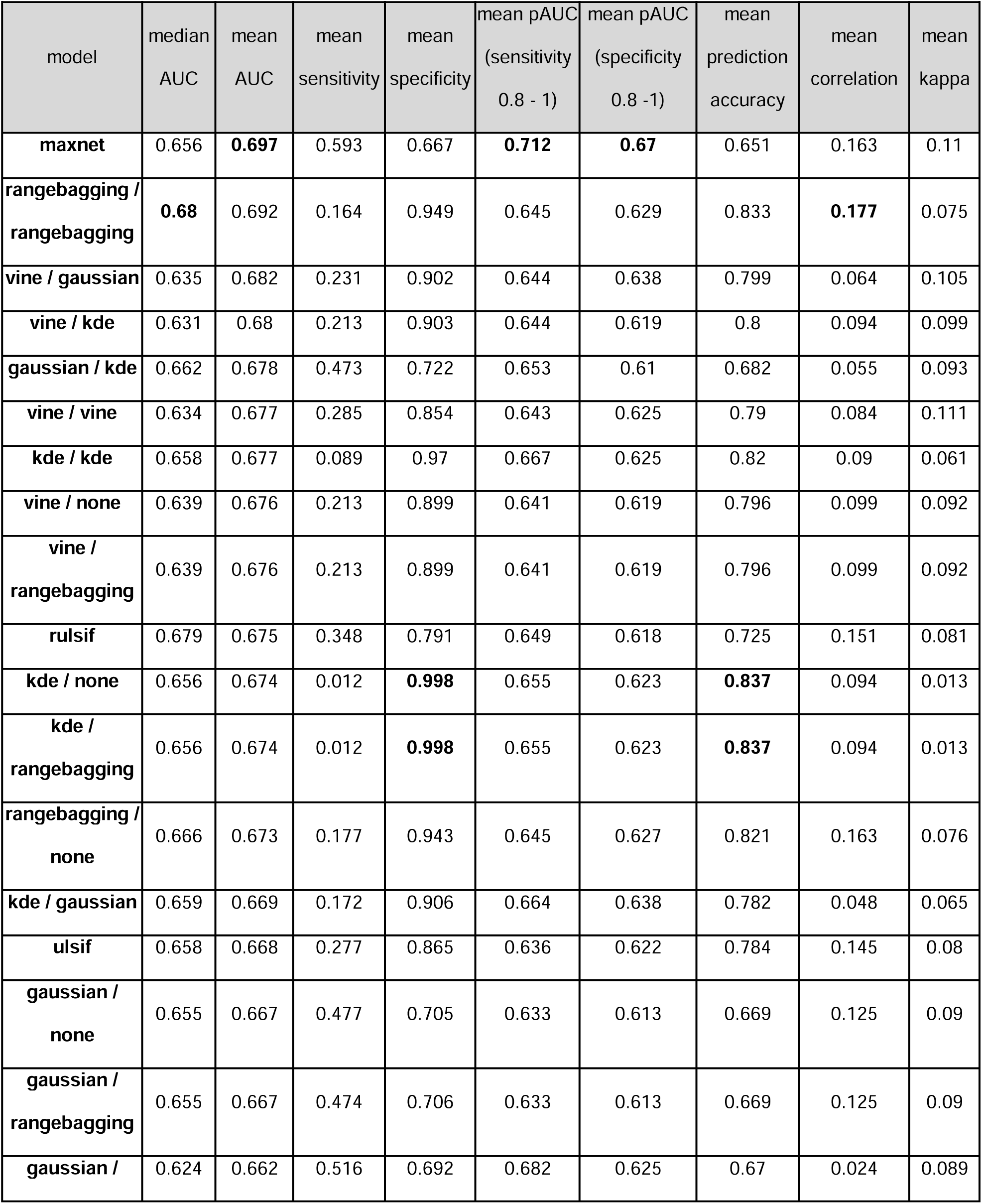
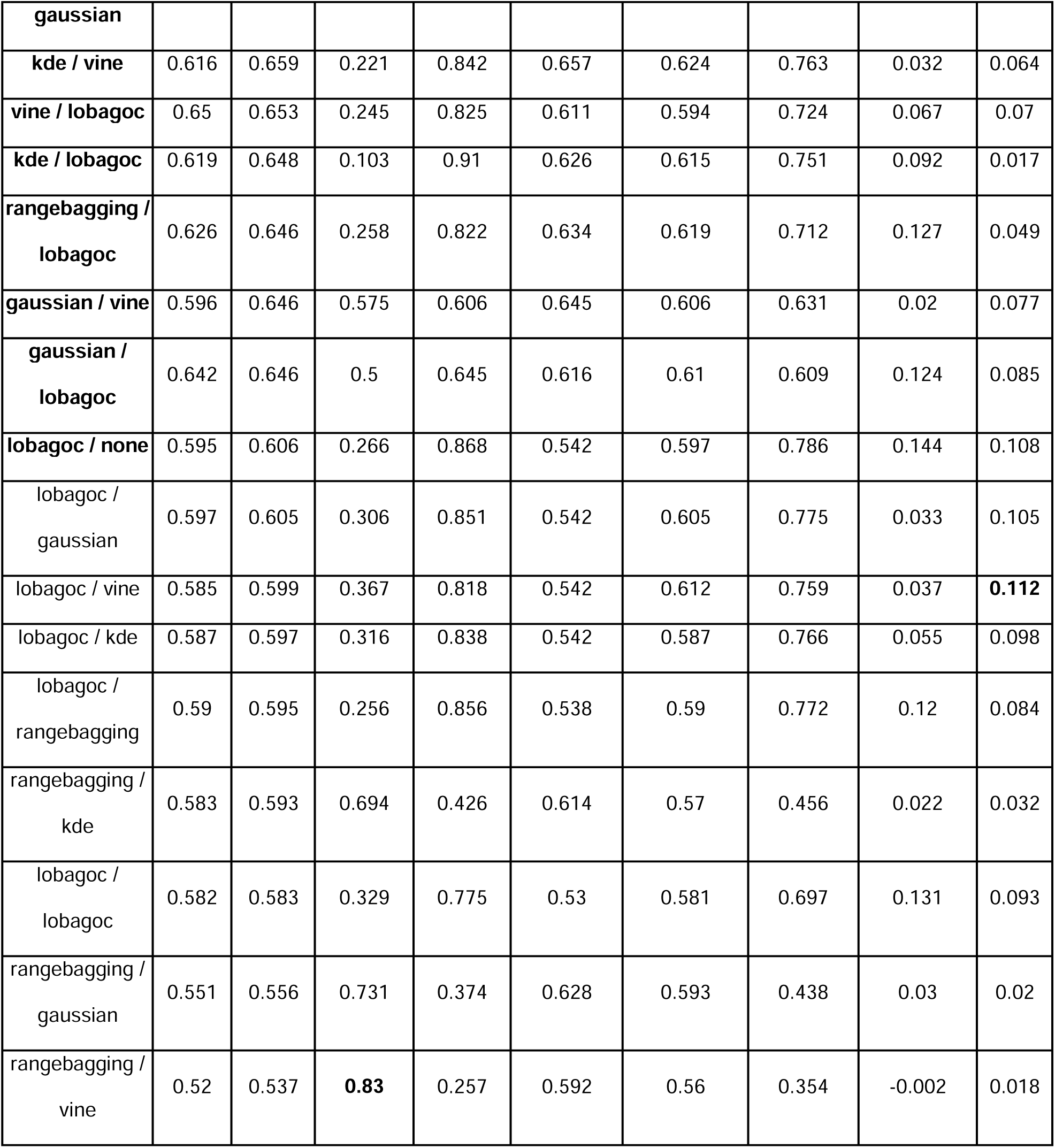
Model performance evaluated with Presence-Absence data, limited to species with 20 or fewer occurrence records. Values shown are the mean values across the entire Elith *et al*. (2020) dataset. AUC refers to the AUC of the full model, where pAUC refers to the partial AUC. Values highlighted in bold are the best performing models for each performance metric. Models in bold have AUCs that are NOT significantly different from Maxnet (Table 3). Range-bagging/range-bagging had the highest median AUC (0.680) while Maxnet models had the highest mean AUCs (0.697). Range-bagging/vine models had the highest mean sensitivity (0.83), but very poor mean AUC scores (0.52). Presence-only KDE models (i.e., KDE / none) had the highest specificity (0.998) and prediction accuracy (0.837). Kappa was highest for the LOBAG-OC/LOBAG-OC models (0.113).

**Table SI 6.**
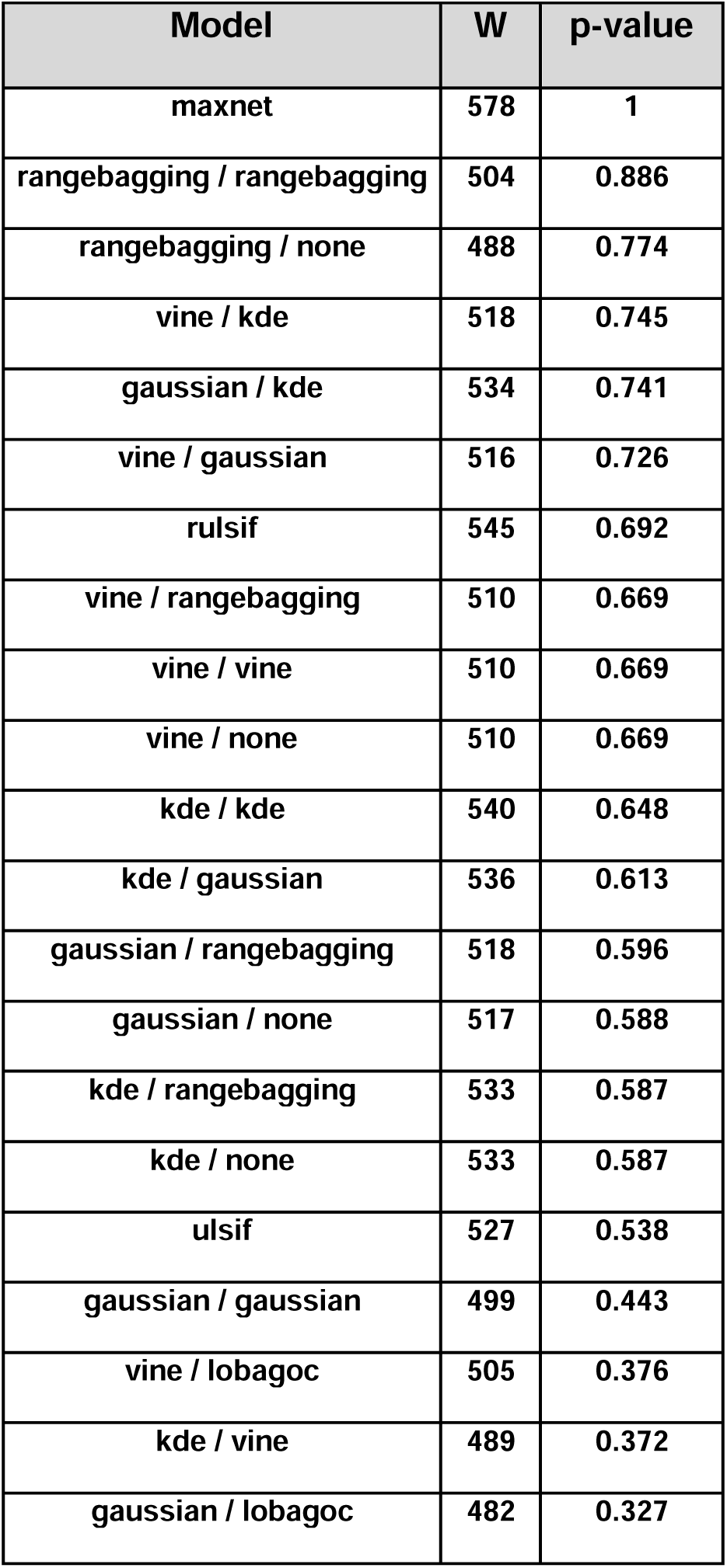
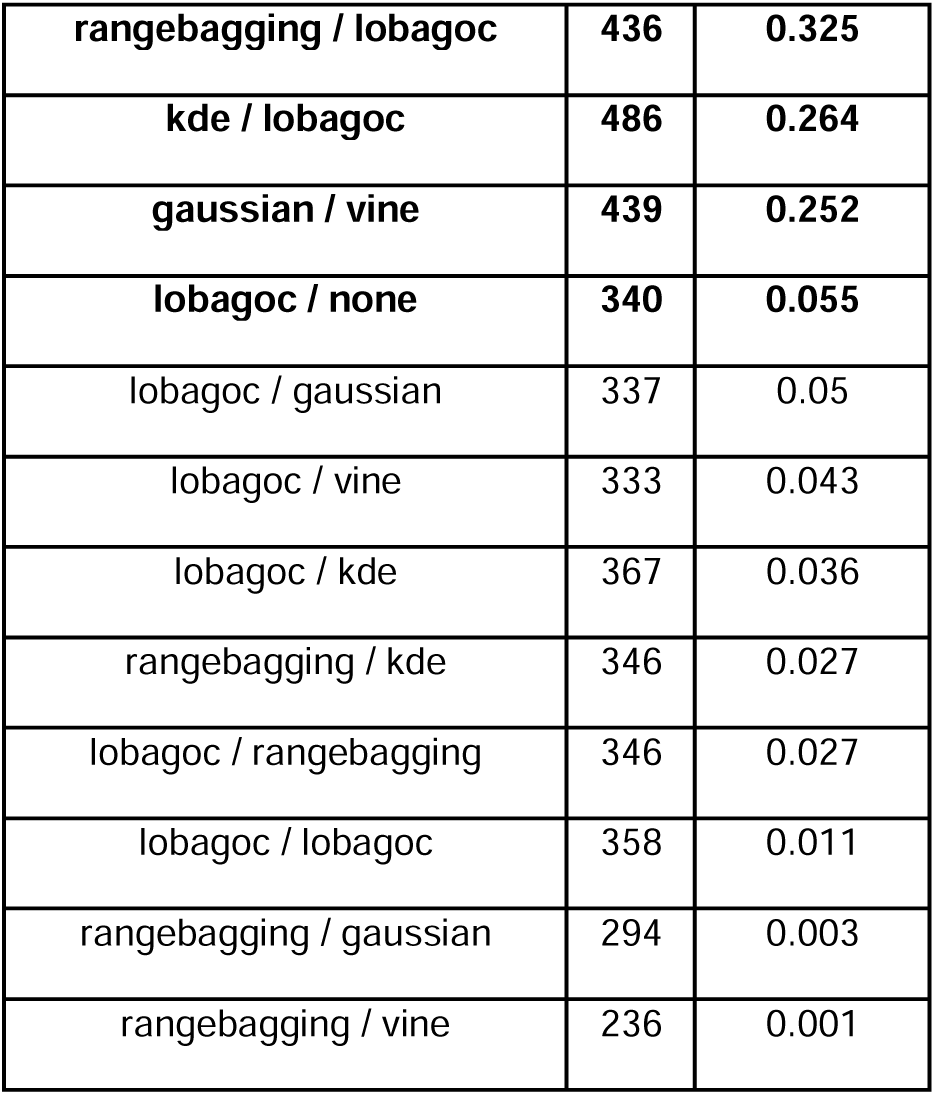
Species with small sample sizes: comparison with Maxnet. Test statistic (W) and p-value refer to the results of a Mann-Whitney test comparing the AUCs of the focal model with those of Maxnet. Comparison was only made for SSSS (those with 20 or fewer records, n = 34). Models in bold are NOT significantly different from Maxnet.

| Model | W | p-value |
| --- | --- | --- |
| maxnet | 578 | 1 |
| rangebagging / rangebagging | 504 | 0.886 |
| rangebagging / none | 488 | 0.774 |
| vine / kde | 518 | 0.745 |
| gaussian / kde | 534 | 0.741 |
| vine / gaussian | 516 | 0.726 |
| rulsif | 545 | 0.692 |
| vine / rangebagging | 510 | 0.669 |
| vine / vine | 510 | 0.669 |
| vine / none | 510 | 0.669 |
| kde / kde | 540 | 0.648 |
| kde / gaussian | 536 | 0.613 |
| gaussian / rangebagging | 518 | 0.596 |
| gaussian / none | 517 | 0.588 |
| kde / rangebagging | 533 | 0.587 |
| kde / none | 533 | 0.587 |
| ulsif | 527 | 0.538 |
| gaussian / gaussian | 499 | 0.443 |
| vine / lobagoc | 505 | 0.376 |
| kde / vine | 489 | 0.372 |
| gaussian / lobagoc | 482 | 0.327 |
| **rangebagging / lobagoc** | **436** | **0.325** |
| **kde / lobagoc** | **486** | **0.264** |
| **gaussian / vine** | **439** | **0.252** |
| **lobagoc / none** | **340** | **0.055** |
| lobagoc / gaussian | 337 | 0.05 |
| lobagoc / vine | 333 | 0.043 |
| lobagoc / kde | 367 | 0.036 |
| rangebagging / kde | 346 | 0.027 |
| lobagoc / rangebagging | 346 | 0.027 |
| lobagoc / lobagoc | 358 | 0.011 |
| rangebagging / gaussian | 294 | 0.003 |
| rangebagging / vine | 236 | 0.001 |

**Figure SI 7.**
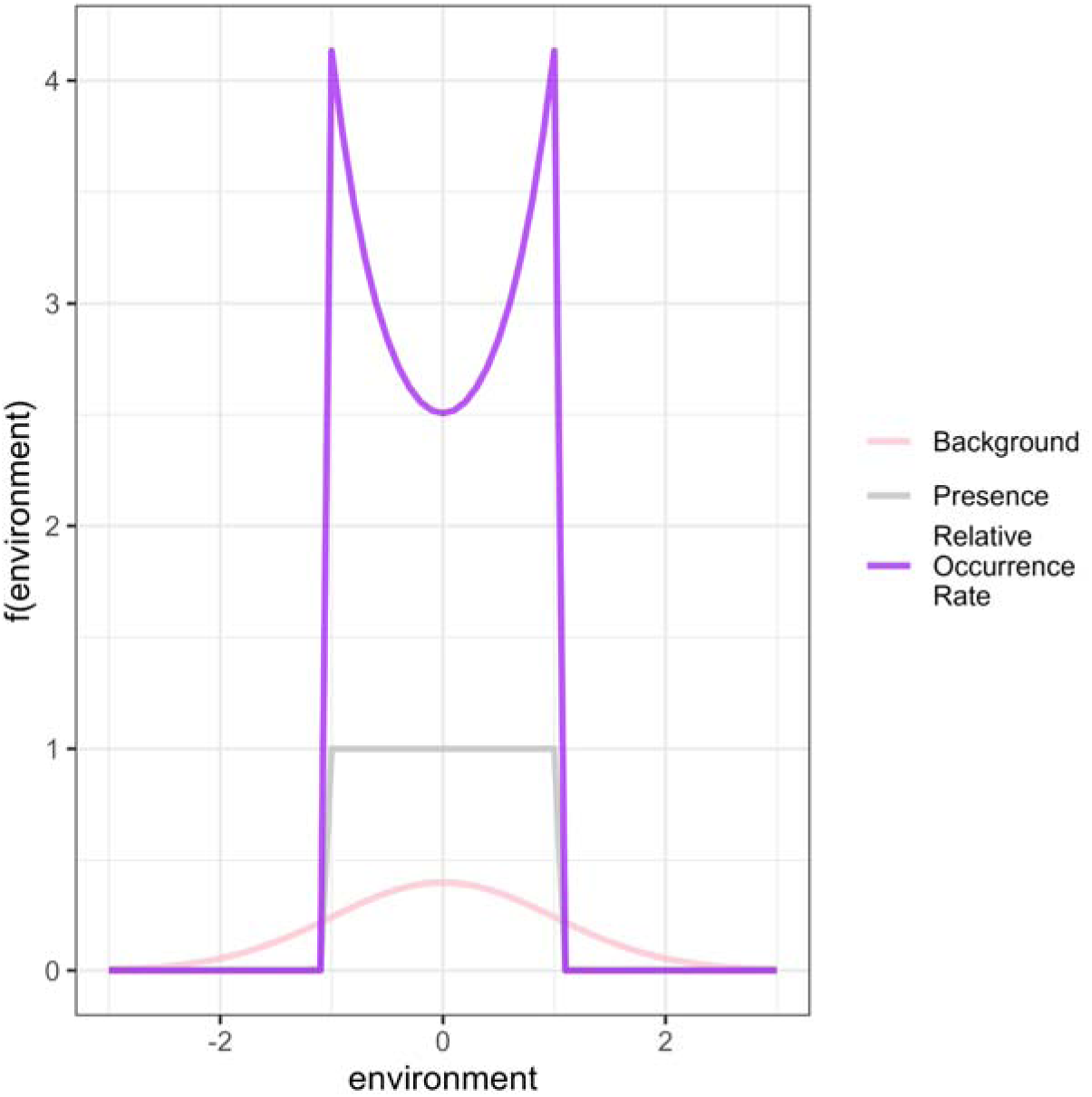
Example showing the shape of a hypothetical range-bagging / Gaussian range model. The presence distribution is modeled with range-bagging estimating in a relatively uniform distribution. The background distribution is gaussian, and the relative occurrence rate is the ratio of the two. This combination of algorithms results in a bimodal distribution. Even though a presence-only rangebagging model or gaussian/gaussian model would include values near zero within a resulting binary range map, the range-bagging/gaussian map may omit them (depending on threshold decisions and training presence locations).

**Figure SI 8.**
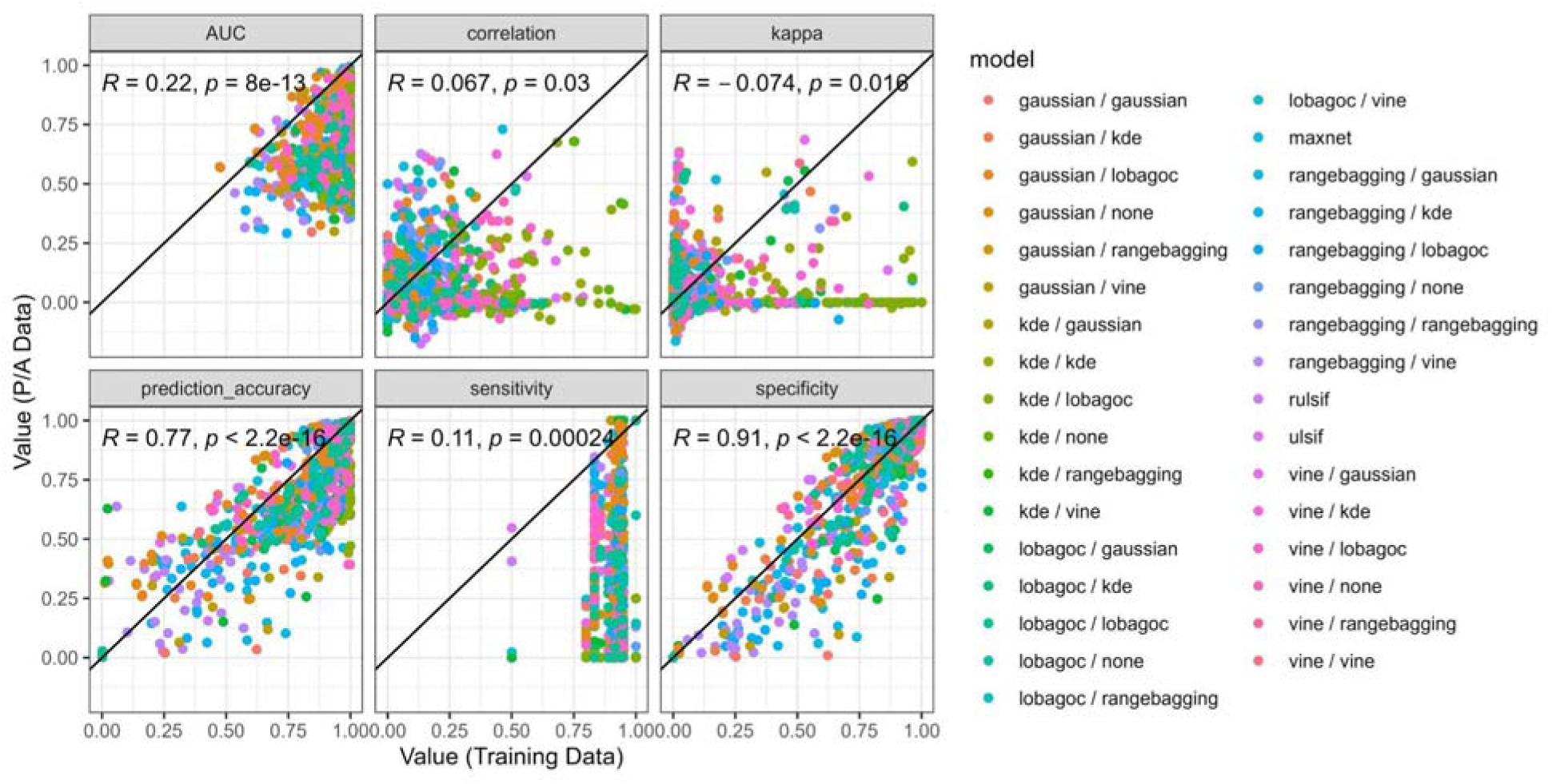
Predictability of model performance for poorly-sampled species. Points represent different species/model combinations. Species shown have been limited to those with 20 or fewer occurrence records in the training data. Lines represent the 1:1 line. Values are Pearson correlation coefficients and corresponding p-values.

**Table SI 9.** Percentages of models for poorly-sampled species for which AUC could not be calculated or was worse than random. The column “Presence/Absence AUC < 0.5” includes both models for which AUC could not be calculated and those where the AUC value was worse than random. Only species with 20 or fewer occurrences were included (n = 34) .

| **Model** | **NAs in Presence/**<br>**Absence AUC**<br>**values (%)** | **Presence/ Absence**<br>**AUCs < 0.5 (%)** |
| --- | --- | --- |
| **rangebagging / rangebagging** | **0** | **7** |
| **gaussian / none** | **0** | **9** |
| **gaussian / rangebagging** | **0** | **9** |
| **vine / gaussian** | **3** | **9** |
| **gaussian / kde** | **0** | **12** |
| **gaussian / lobagoc** | **0** | **12** |
| **kde / none** | **0** | **12** |
| **kde / rangebagging** | **0** | **12** |
| **rulsif** | **0** | **12** |
| **ulsif** | **0** | **12** |
| **vine / kde** | **3** | **12** |
| **vine / none** | **3** | **12** |
| **vine / rangebagging** | **3** | **12** |
| **vine / vine** | **3** | **12** |
| **rangebagging / lobagoc** | **0** | **13** |
| **rangebagging / none** | **0** | **13** |
| **kde / kde** | **0** | **15** |
| **kde / lobagoc** | **0** | **15** |
| **maxnet** | **0** | **15** |
| **vine / lobagoc** | **0** | **15** |
| **gaussian / gaussian** | **0** | **18** |
| kde / gaussian | 0 | 21 |
| kde / vine | 3 | 21 |
| rangebagging / kde | 0 | 23 |
| gaussian / vine | 6 | 24 |
| lobagoc / kde | 9 | 26 |
| lobagoc / lobagoc | 3 | 26 |
| lobagoc / gaussian | 18 | 32 |
| lobagoc / rangebagging | 12 | 32 |
| lobagoc / vine | 18 | 32 |
| lobagoc / none | 18 | 35 |
| rangebagging / gaussian | 0 | 37 |
| rangebagging / vine | 7 | 50 |

**Table SI 10.** Ensemble performance relative to component models. Mean performance comparison for ensemble approaches and component models.

| model | prediction accuracy | specificity | sensitivity | kappa |
| --- | --- | --- | --- | --- |
| ensemble unanimous<br>support | 0.825 | 0.979 | 0.064 | 0.046 |
| KDE/KDE | 0.82 | 0.97 | 0.089 | 0.061 |
| ensemble mean | 0.741 | 0.852 | 0.247 | 0.056 |
| ruLSIF | 0.725 | 0.791 | 0.348 | 0.081 |
| **Maxnet** | 0.645 | 0.658 | 0.599 | **0.109** |
| **ensemble any**<br>**support** | 0.622 | 0.608 | **0.648** | 0.099 |

**Figure SI 11.**
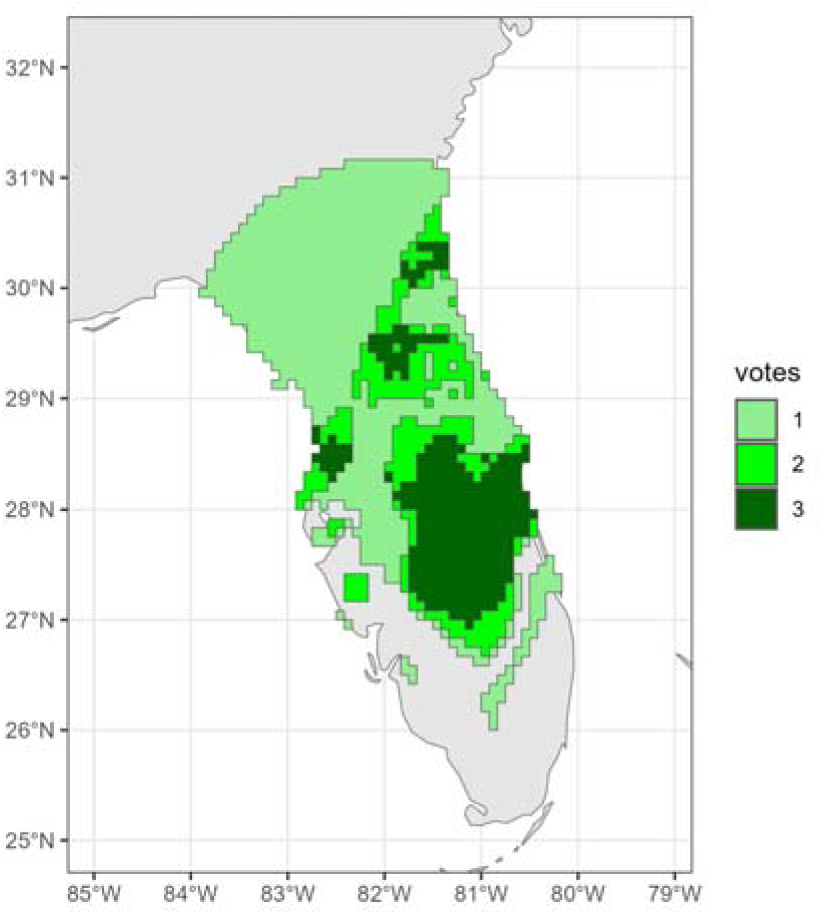
Example ensemble prediction for *Chrysopsis subulata*, a species with 15 validated observations in the BIEN dataset. Data from BIEN (B. S. Maitner et al., 2017).

